# Under pressure: Evolutionary trade-offs shape the single visual opsin of deep-sea octopods

**DOI:** 10.64898/2026.07.15.738692

**Authors:** Giacinto De Vivo, Minglei Ma, Giobbe Forni, Andrea Luchetti, Fabio Crocetta, Marjorie A. Liénard, Salvatore D’Aniello

## Abstract

Octopods possess remarkable camera-type eyes and specialised image-forming vision that support orientation, prey detection, predator avoidance and visual communication. Unlike vertebrates and arthropods, however, octopod vision is thought to rely mainly on a single rhodopsin (r-opsin1), raising the question of how such a constrained system adapts across contrasting light environments from shallow coastal waters to deep-sea habitats.

Using transcriptional profiling of the retina and optic lobe from seven native octopod species of the Gulf of Naples, we show that r-opsin1 is the predominant visual gene across all species and investigate how contrasting photic habitats have shaped its molecular and functional evolution. Although positive selection analyses revealed no general association between habitat depth and r-opsin1 evolution, twelve codons in the deep-mesopelagic *Pteroctopus tetracirrhus* r-opsin1 showed evidence of positive selection, including two residues located on opposing helices of the retinal-binding pocket.

*In vitro* experiments demonstrated that substituting either I87V and F201N produced a marked bathochromatic shift from the wild-type blue-green spectrum towards red wavelengths, whereas their combination restored the wild-type spectral profile. Other deep-sea octopod r-opsin1 naturally bearing one of these substitutions retained blue-green sensitivity. Reconstructed ancestral proteins similarly maintained blue-green absorption, supporting conservation of this spectral phenotype throughout octopod r-opsin1 evolution.

We further show that F201N reduces adiabatic compressibility within the retinal-binding pocket, by co-evolving compensatory sites offering a structural trade-off, favouring pressure adaptation while restraining spectral tuning. Together, our findings support that octopod visual rhodopsin evolution has been shaped by the multiple ecological pressures of contrasting marine environments, illustrated with lineage-specific and habitat-dependent trajectories on a single locus while preserving a conserved visual phenotype.

## Introduction

Octopods are among the most distinctive cephalopods, a group that also includes squid, cuttlefish and nautiluses, attracting considerable interest in their advanced cognition and complex behaviours. Over their evolutionary history, the different octopod species colonized a wide range of habitats, including coastal, benthic, pelagic, and deep-sea environments (Albertin et al., 2015; Destanović et al., 2023; Hanke and Kelber, 2020; Yoshida et al., 2015; Young, 1991). Like other extant cephalopods, they are active predators that rely on multiple sensory modalities, as well as remarkable camouflage abilities, to capture prey and avoid predators. In this animal group, vision plays an important role, as evidenced by highly developed camera-type eyes equipped with a lens, iris, cornea, retina, and the optic lobes, which represent two-thirds of their brain and are entirely dedicated to visual processing (Hanke and Kelber, 2020; Yoshida et al., 2015). The structure of their retina, with the rhabdomeres of photoreceptors oriented perpendicular to each other, makes them able to perceive polarized light. Cephalopods rely on image-forming vision not only to detect prey or to avoid predators but also to communicate and orient themselves (Hanke and Kelber, 2020).

Beyond the morphology of the visual system, vision depends on a specialized molecular machinery for light detection, which relies on the opsin signalling cascade. Opsins are the primary photoreceptive molecules underlying animal vision and mediate a variety of non-visual light-dependent processes, including circadian entrainment, seasonal regulation, and orientation in the water column (Davies et al., 2010; Gühmann et al., 2015; Terakita, 2005). They belong to the G-protein-coupled receptor (GPCR) family and contain seven transmembrane domains that covalently bind a vitamin A-derived chromophore, the retinal, via a protonated Schiff base to a conserved lysine residue. In bovine rhodopsin, this residue corresponds to position 296 in the amino acid sequence (Lys-296). This interaction confers light sensitivity: in darkness, the retinal is bound in the 11-*cis* configuration, whereas photon absorption induces isomerization to the all-*trans* isomer, forming the photoactive state (Figure 1A,B). The resulting conformational change activates the opsin and triggers the associated G-protein signalling cascade (Terakita, 2005). Opsins can be monostable or bistable based on the stability of their photoactive state. Whereas most vertebrate monostable opsins release retinal following photohydrolysis (bleaching), certain invertebrate opsins are bistable and retain a covalent linkage to retinal throughout the photoreaction cycle (Palczewski et al., 2000; Terakita, 2005). In these opsins, the 11-*cis* configuration can be regenerated upon absorption of a second photon, restoring the inactive state without bleaching (Tejero et al., 2024). Intriguingly, despite the complexity of their visual system, cephalopods rely on a reduced number of opsins compared to vertebrates and arthropods (De Vivo et al., 2023), achieving sophisticated image-forming vision through a single visual opsin.

**Figure 1.**
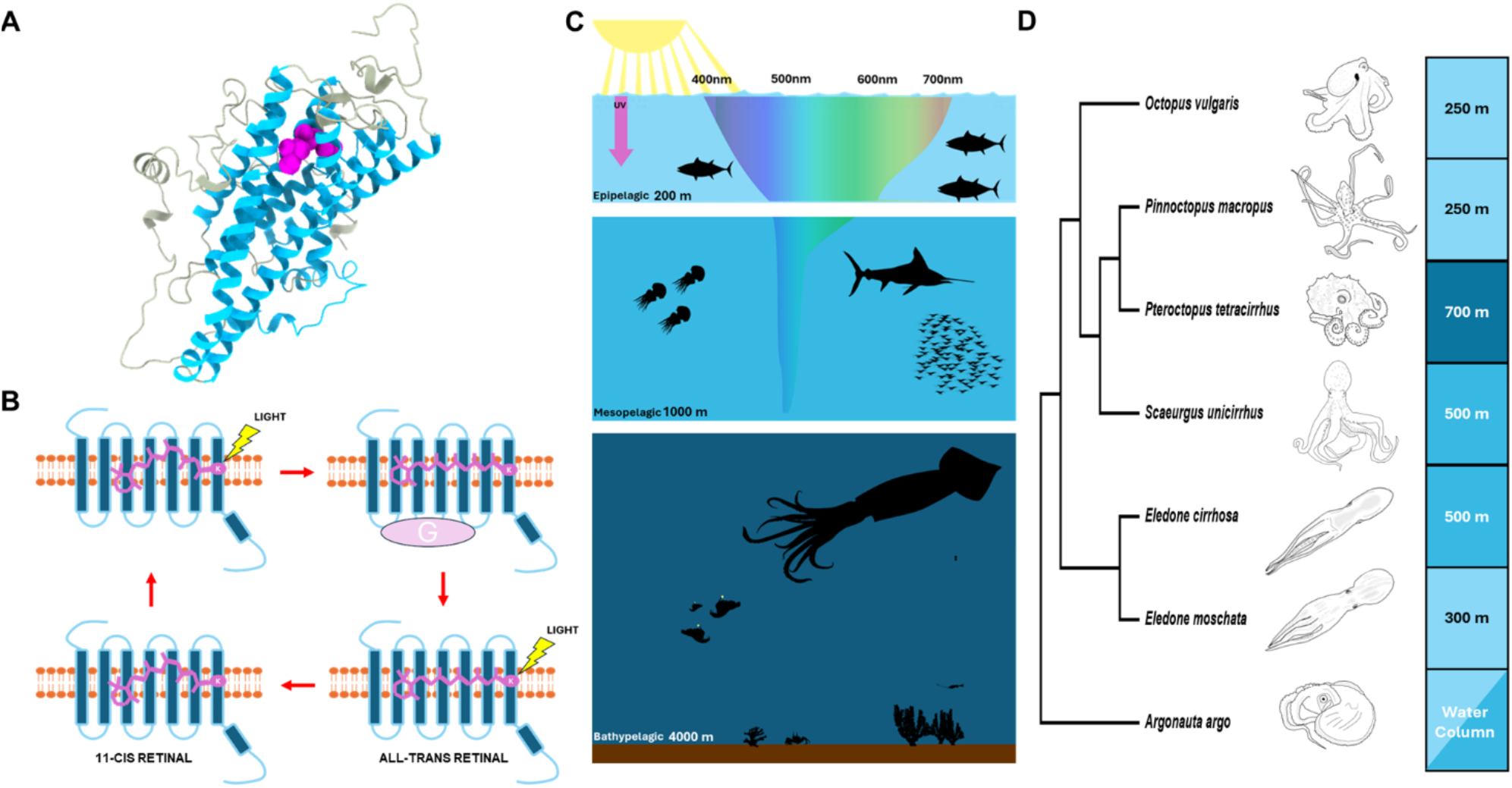
Visual ecology of octopod r-opsin1. (A) Three-dimensional structure of r-opsin1 of *Octopus vulgaris* showing the transmembrane (TM) helices in celestial blue and the retinal chromophore in purple. The 11-*cis*-retinal is covalently attached to Lys305 (corresponding to Lys296 in bovine rhodopsin) on helix VII. (B) Schematics of the proposed visual cycle of cephalopods’ r-opsin1: upon photon absorption, retinal changes from 11-*cis* to all-*trans*, inducing a conformational change in the opsin that allows it to bind the G-protein and initiate the signalling cascade. A second photon interaction can revert retinal from all-*trans* back to 11-*cis*. (C) In marine photic environments, sunlight undergoes attenuation in the water column: red and violet wavelengths are scattered and absorbed at depths below 200 m, whereas blue-green light reaches the mesopelagic zone up to 1000 m below the surface. (D) Phylogenetic relationships between the octopod species of the Gulf of Naples’ ecosystem, including their typical maximum habitat depth. The phylogeny is adapted after López-Córdova et al. (2022), Sánchez-Márquez et al. (2022) and Taite et al. (2023) with maximum habitat depth traits mapped from Jereb et al. (2005).

Opsin molecular adaptations to different light conditions in nature are well documented in deep-sea fishes, cetaceans and nocturnal insects (Feuda et al., 2016; Hagen et al., 2023; Musilova et al., 2019; Porter et al., 2016; Ricci et al., 2022; Ricci et al., 2023). The evolutionary success of these adaptations depends on coping with four main factors: drastic decrease in light intensity, increased scattering and absorption of red and violet wavelengths (leaving only blue-green light around 480 nm below ∼200 m), bioluminescence presence, and high hydrostatic pressure, which may impact protein function (Figure 1C).

Vertebrates and arthropods typically use multiple opsin orthologues for vision, allowing them to discriminate among different colours in both terrestrial and marine environments (De Vivo et al., 2023; Jacobs, 2009; Liénard et al., 2021; Musilova et al., 2019; Ricci et al., 2022; Ricci et al., 2023; Roberts et al., 2025). This photoreceptor richness has subsequently been re-adapted in deep-sea species to cope with the extreme light reduction. For instance, many deep-sea fishes have undergone major molecular adaptations, losing red- and violet-sensitive opsins while expanding their green- and blue-sensitive opsin toolkit through gene duplications and subsequent functional diversification (Musilova et al., 2019). In contrast, cetaceans and bats exhibit a reduced complement of opsins, generally retaining rod opsin (RH1) and mid- to long-wavelength–sensitive opsins (MWS/LWS), while the short-wavelength–sensitive opsin (SWS1) is pseudogenized (Simões et al., 2018; Meredith et al., 2013). In addition, cetaceans that dive into the deep sea, such as sperm whales, have lost MWS/LWS and rely exclusively on RH1, becoming rod monochromats. Importantly, opsin pseudogenization events, as well as the evolution of monochromacy in some cetacean species, are associated with secondary loss and, in some cases, may represent trade-offs between vision and the evolution of echolocation in these groups (Thiagavel et al., 2018). Despite echolocation playing a key role in some cetaceans, opsins show signs of positive selection that led to blue shifted sensitivity associated with diving behaviours (Dungan et al., 2016; Dungan and Chang, 2022).

In cephalopods, the situation is different. They are thought to rely primarily on a single visual rhabdomeric opsin, named octopus rhodopsin (hereby r-opsin1) (Bonadè et al., 2020; Chung and Marshall, 2016; Hanke and Kelber, 2020; Hara et al., 1967; Hara & Hara, 1972; Yoshida et al., 2015), supporting monochromatic vision that is not the result of secondary loss, but rather a trait shared by all cephalopods, likely already present in their common ancestor. Therefore, monochromacy is likely a feature retained throughout cephalopods’ evolutionary history despite the complexity of their visual system and the central importance that vision has for these animals when compared with other senses (Messenger et al., 1973; Murakami and Kouyama, 2008). How such a constrained genetic system adapts at the molecular level across highly contrasting environments spanning bright and dim light is intriguing from an evolutionary perspective.

Cephalopod r-opsin1 gene is a Gq-coupled opsin homolog to many invertebrate visual opsins, including those of arthropods (Murakami and Kouyama, 2008). Unlike vertebrate visual c-opsins which are monostable Gt-coupled opsins, squid r-opsin1 has been shown to be bistable (Koyanagi et al 2008; Terakita, 2014). While the structural and biophysical properties of squid r-opsin1 have been extensively studied (Hubbard and St. George, 1958; Ota et al., 2006; Murakami and Kouyama, 2008), the range of molecular evolution and functional variation across opsins in deep-sea cephalopod groups remain unresolved. Earlier studies suggested that deep-sea Decapodiformes (squid, cuttlefish, bobtail) exhibit a blue shift in spectral sensitivity compared to littoral species (Chung and Marshall, 2016). Similar patterns have been reported in other taxa inhabiting environments with marked differences in light conditions, such as deep-sea fish and cetaceans, where specific amino acid substitutions have been shown to play key roles in spectral tuning (Musilova et al., 2019; reviewed in Hagen et al., 2023;). However, none of the octopod species examined in Chung and Marshall (2016), or in other studies on opsin spectral tuning, spend most of their life cycle in the deep sea (200-1000m). Therefore, our understanding of octopod r-opsin1 evolution and adaptation to dim light environments remains incomplete (Fig. 1C).

Intrigued by the extreme versatility of high- and low-light habitats that octopods have colonised despite the conserved single molecule-based Swiss-army-knife-like photovisual system, we set out to investigate the functional diversity and selection regimes acting on r-opsin1 associated with octopod species from distinct light environments and sea-depth habitats. For this, we focused on the Gulf of Naples as our marine ecosystem (Italy, Western-Central Mediterranean Sea), an internationally recognized biodiversity hotspot harbouring seven incirrate octopods spanning several ecological niches, from littoral to deep-sea environments (Figure 1C,D), but also distinct positions within these habitats - benthic *i.e.* near the seafloor and pelagic *i.e* in the water column. To explore the role of environmental factors, such as habitat depth, light intensity, spectral quality, and hydrostatic pressure as potential evolutionary drivers shaping octopod visual systems, we integrated transcriptomics, molecular evolutionary analyses, structural modelling, functional assays and phylogenomic analyses. We investigate how a visual system constrained to a single-visual opsin evolved across marine light gradient habitats, and if ecological diversification required repeated changes in spectral sensitivity or could be maintained through alternative molecular mechanisms.

## Results

### Positive evolutionary selection acts on r-opsin1 in octopods

Our study system, the Gulf of Naples, supports a wide range of marine habitats, from coastal seagrass meadows and rocky reefs to deep-sea canyons and pelagic habitats, offering controlled geographical parameters such as annual turbidity and latitude, as well as robust ecological and geological historical record, thereby forming a unique ecosystem for studying the correlated evolution of marine ecosystems, assemblages, and species. Field sampling collection of native octopus occurred exclusively in the Gulf of Naples ensuring known geographical conditions and, therefore, a correlation between depth and light conditions in our comparative analyses. Fresh live bycatch octopus’ specimens were obtained from local fishermen prior to careful tissue collection and RNA-sequencing of the visual system, comprised of the retina and optic lobe tissues. The following incirrate octopods were selected including two species mostly epipelagic, namely *Octopus vulgaris* (common octopus) and *Pinnoctopus macropus* (white-spotted octopus), and four mostly mesopelagic species, namely *Pteroctopus tetracirrhus* (fourhorn octopus), *Scaeurgus unicirrhus* (unihorn octopus), *Eledone moschata* (musky octopus) and *Eledone cirrhosa* (curled octopus), whereas the pelagic octopus *Argonauta argo* was included as a phylogenetic and ecological outgroup. Transcriptomic profiling of the retina and optic lobe for *P. macropus*, *P. tetracirrhus*, *S. unicirrhus*, *E. moschata* and *E. cirrhosa* retrieved expression data for six photosensitive-like transcripts, namely r-opsin1 (also earlier referred to as rhodopsin), r-opsin2, xenopsin, retinochrome, peropsin, and pseudospin (Figure S1). Overall, there is generally low expression of xenopsin, r-opsin2, peropsin, and pseudopsin transcripts in the visual system across octopus, whereas the r-opsin1 octopod rhodopsin appears to be highly expressed. Hence, the log_2_ fold expression of r-opsin1 is almost double the expression of retinochrome, the second most expressed opsin, across all species analyzed (Figure S1), supporting a central role in octopod photoreception.

We next investigated whether a specific selective regime acts on r-opsin1 in deep-sea species, as this could indicate constraints or adaptations associated with low-light environments. For this, we computed a species tree using r-opsins from the different species obtaining topology is consistent with known species phylogeny (López-Córdova et al., 2022; Sánchez-Márquez et al., 2022; Taite et al., 2023) and subsequently implemented a codon-based phylogenetic likelihood approach to test whether specific genes or codon sites evolved under selective pressure favouring beneficial mutations (Álvarez-Carretero et al., 2023). This approach assumes that molecules under selection show a statistically distinct ratio of non-synonymous to synonymous substitutions (dN/dS). We tested several scenarios against the null hypothesis (M0), which assumes the same dN/dS ratio across all sites and lineages, to assess whether the mode of evolution of r-opsin1 differs between littoral and deep-sea species (Table 1, File S1). In particular, the first model (M1), which allows each branch to have a distinct dN/dS ratio, indicated significant differences from M0, suggesting heterogeneity in selective pressure among branches. We then tested which branches were responsible for the differences in dN/dS identified by M1. Specifically, we applied a second model (M2), in which predefined groups are allowed to have different dN/dS ratios, to two alternative hypotheses: (i) littoral species are under positive selection, or (ii) deep-sea species are under positive selection. Our results showed no evidence of distinct selective regimes associated with habitat type. Likewise, no codon sites under selection were identified using the branch-site model (BSM) when deep-sea and littoral groups were compared in either direction.

**Table 1.**
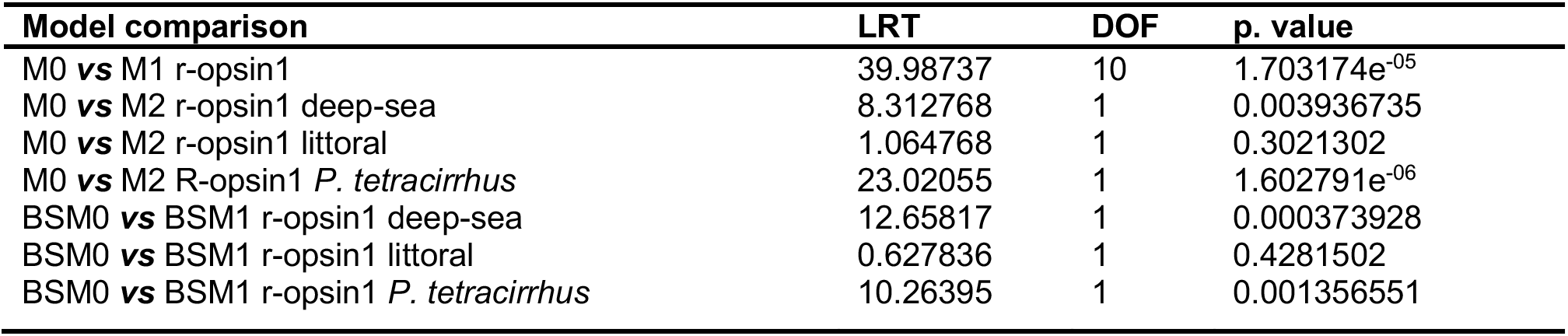
Comparison between different positive selection models on r-opsin1 in PAML. LRT indicates Likelihood Ratio Test, and DOF indicates the difference in Degrees Of Freedom. *Argonauta argo* was used as the outgroup for the analysis.

| Model comparison | LRT | DOF | p. value |
| --- | --- | --- | --- |
| M0 vs M1 r-opsin1 | 39.98737 | 10 | 1.703174e<sup>-05</sup> |
| M0 vs M2 r-opsin1 deep-sea | 8.312768 | 1 | 0.003936735 |
| M0 vs M2 r-opsin1 littoral | 1.064768 | 1 | 0.3021302 |
| M0 vs M2 R-opsin1 *P. tetracirrhus* | 23.02055 | 1 | 1.602791e<sup>-06</sup> |
| BSM0 vs BSM1 r-opsin1 deep-sea | 12.65817 | 1 | 0.000373928 |
| BSM0 vs BSM1 r-opsin1 littoral | 0.627836 | 1 | 0.4281502 |
| BSM0 vs BSM1 r-opsin1 *P. tetracirrhus* | 10.26395 | 1 | 0.001356551 |

We then applied M2 to test whether selection might act on individual species rather than on habitat-defined groups, thereby explaining the results observed under M1. This analysis revealed that one r-opsin1 sequence, belonging to *P. tetracirrhus* (hereafter referred to as PteRh1), shows clear signatures of positive selection. Specifically, under the M1 model, the dN/dS ratio estimated for PteRh1 was 0.451702, whereas all other r-opsin1 sequences showed much lower values, consistent with strong functional constraint: 0.0829095 in *O. vulgaris*, 0.0228 in *P. macropus*, 0.0422 in *S. unicirrhus*, 0.0779 in *E. moschata*, 0.1280 in *E. cirrhosa*, and 0.0849 in *A. argo*. Similarly, the M2 branch model estimated a foreground dN/dS ratio of 0.4457 for *P. tetracirrhus*, compared with 0.0803 for the background branches. Together with the higher likelihood ratio test (LRT) value and lower p-value relative to alternative models grouping species by habitat, these results suggest that, although purifying selection acts overall on r-opsin1, a relaxation of functional constraints may have acted preferentially along the evolutionary branch leading to the *Pteroctopus* lineage. To double check these results we performed a second independent positive selection analysis, aBSREL, using HyPhy v2.6 (Smith et al., 2015). This analysis confirmed the PAML analysis (File S1).

Subsequently, to identify which sites are under selection in *P. tetracirrhus*, we applied a branch-site model with Bayes Empirical Bayes (BEB) inference, designating *P. tetracirrhus* as the foreground lineage. This analysis detected 12 codon sites in PteRh1 with a high posterior probability > 0.8 of being under positive selection (Figure 2), including four sites with strong statistical support (posterior probability > 0.95).

**Figure 2.**
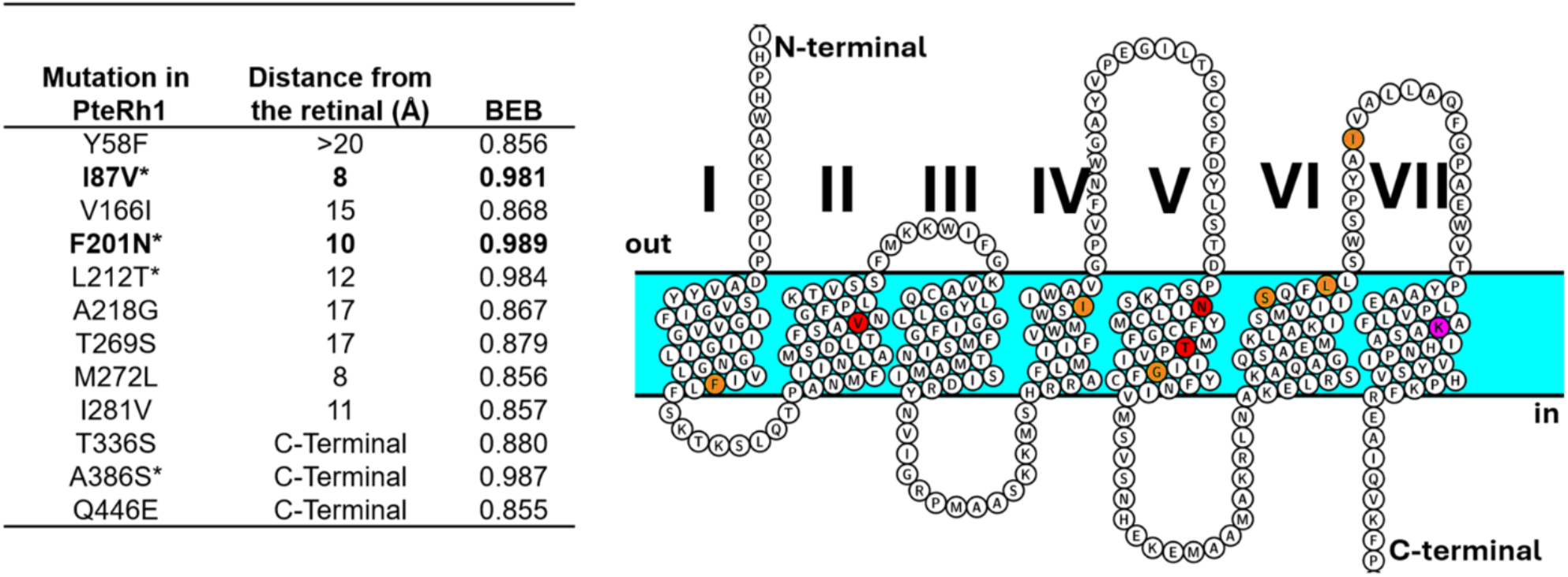
Residues under selection in *Pteroctopus tetracirrhus* r-opsin1. The table shows positively selected mutations found in *P. tetracirrhus* but not in other octopods species from the Gulf of Naples, the distance of each residue relative to any atom of the retinal binding pocket and the BEB probability. The snake plot shows the residue position within the transmembrane domains (TM) of primary protein sequence. The * and the red indicates residues with BEB > 0.95, the orange indicates residues with BEB < 0.95 and >0.8; the purple indicates the retinal binding domain (Lys305, corresponding to Lys296 in bovine rhodopsin). Residues are numbered according to the amino acid positions in PteRh1. The 3 C-terminal residues are not shown on the snake plot.

### Selected substitutions have a compensatory effect on r-opsin1 sensitivity to light

The indication of positive selection of many codons within PteRh1 inspired us to verify where they are positioned within the receptor, which first called for obtaining a reliable homology structure, which we generated using an iterative Threading ASSEmbly Refinement (I-TASSER) structure prediction (Zheng et al., 2025) (Figure 3; File S2). Of all twelve amino acids under positive selection none appear homologous to sites previously identified in cephalopod as adaptation to higher hydrostatic pressure (Porter et al., 2016), and five residues were found to lie proximal (<12 Å) to the retinal chromophore (Figure 2, Figure 3). Among all variant residues under positive selection in *P. tetracirrhus*, we focused on I87V and F201N, based on their posterior probability (BEB), distance with known important opsin residues, and orientation relative to the retinal in the modelled 3-D structure (Figure 3). Our modelling analysis indicates that residue 87 is positioned 3 to 7 Å from residues lining the retinal binding pocket, namely F84, N88, and Y112, whereas 201 is localized within 6 Å of F189 and M205, two residues predicted to be involved in Schiff base stabilization, efficient photoactivation, and postulated to affect spectral tuning (Chung and Marshall, 2016). The modelling also indicates that residues 87 and 201 are localised on helices placed on opposite sides of the binding pocket.

**Figure 3.**
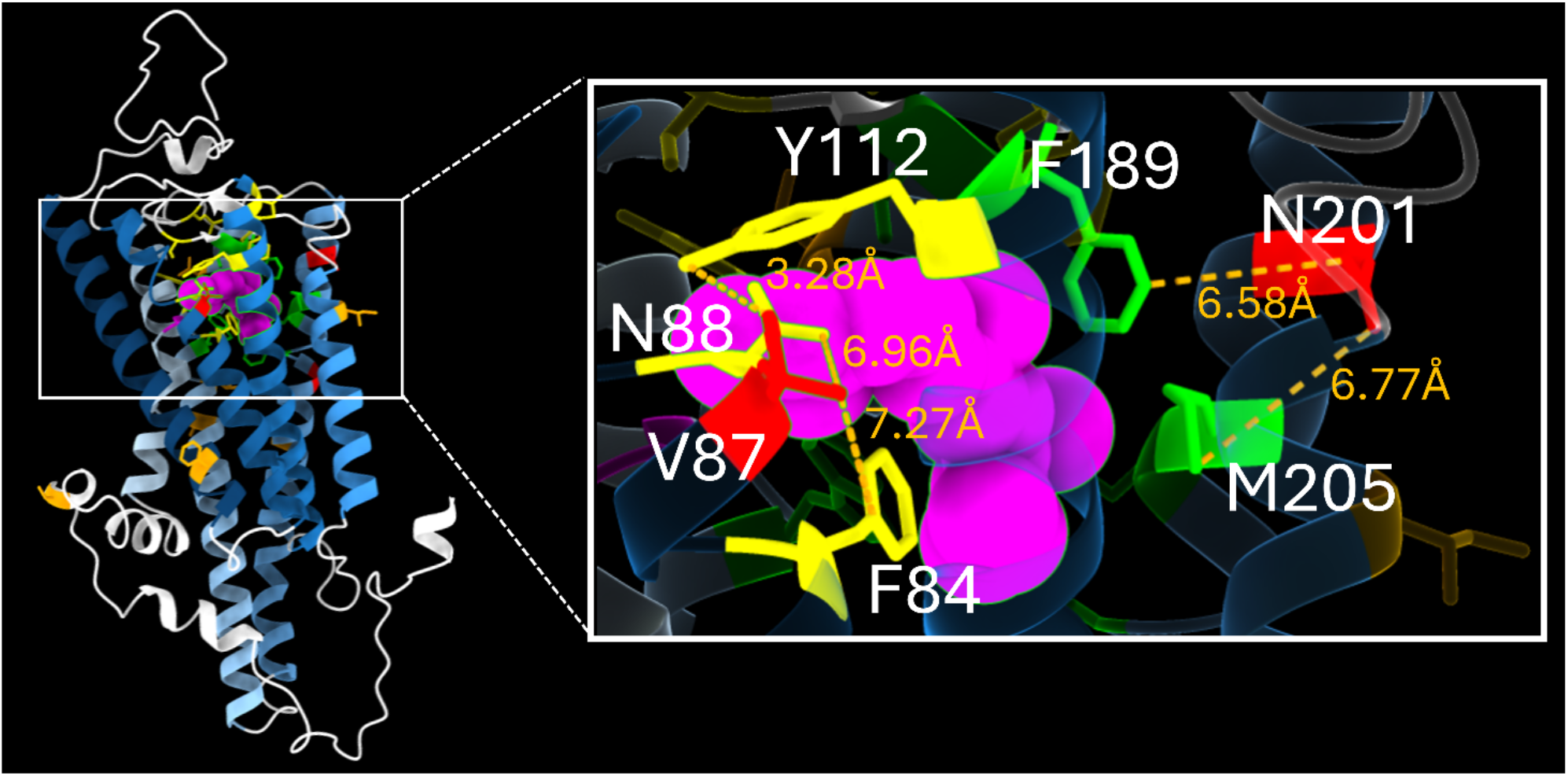
Interactions of sites under selection in *Pteroctopus tetracirrhus* r-opsin1. Protein structure of PteRh1 reconstructed using I-Tasser showing the molecule amino-acidic sites from 1 to 378. The zoom section shows distances between V87 (Helix 2) or N201 (Helix 5) and the residues in the chromophore binding pocket (dashed lines). Colours are as follows: Transmembrane helices are in light blue, the retinal chromophore and K296 Schiff base complex are in purple, sites with BEB > 0.9 are in red, sites with 0.8 < BEB ≤ 0.95 are in orange, residues predicted to be involved in retinal isomerization are in yellow, and residues predicted to be associated with cephalopod spectral tuning in green (Chung and Marshall, 2016), blue-celestial indicate the different TM domains. Residues are numbered according to the amino acid positions in PteRh1. All mutations are numbered according to the amino acid positions in PteRh1. To facilitate comparison with other rhodopsin models, an alignment with homologous positions in well-studied opsins, including squid, jumping spider JSR1, and bovine rhodopsins as reference, is also provided in File S3.

To determine whether these differences in amino acidic composition result in changes to the λ_max_, we first predicted the absorption spectra of the molecules under investigation by initially performing an *in silico* analysis using VPOD (Table S1), which suggested no major differences in the λ_max_ values of PteRh1, OvuRh1, or other examined octopod Rh1. Prior to test a potential role for residues 87 and 201, we first established a functional baseline and measured the absorption spectra of a wild-type r-opsin1 in the littoral octopod *O. vulgaris* OvuRh1, which carries I87 and F201 similarly to the six other octopod Rh1 opsins investigated, compared to the deep-sea *P. tetracirrhus* PteRh1 bearing V87 and N201 instead. After heterologous expression and purification of full-length rhodopsins in HEK cells in the presence of 11-*cis*-retinal, both reconstituted visual pigments absorbed wavelength light ranging from 450 nm to 550 nm with a maximal absorption (λ_max_) at 481 nm ± 4.6 nm (Figure 4A,B, Figure S2A), which is in line with earlier measurements from retinal extracts for *O. vulgaris* (Chung and Marshall, 2016). This result indicates no difference in light absorption between the two species associated with depth.

**Figure 4.**
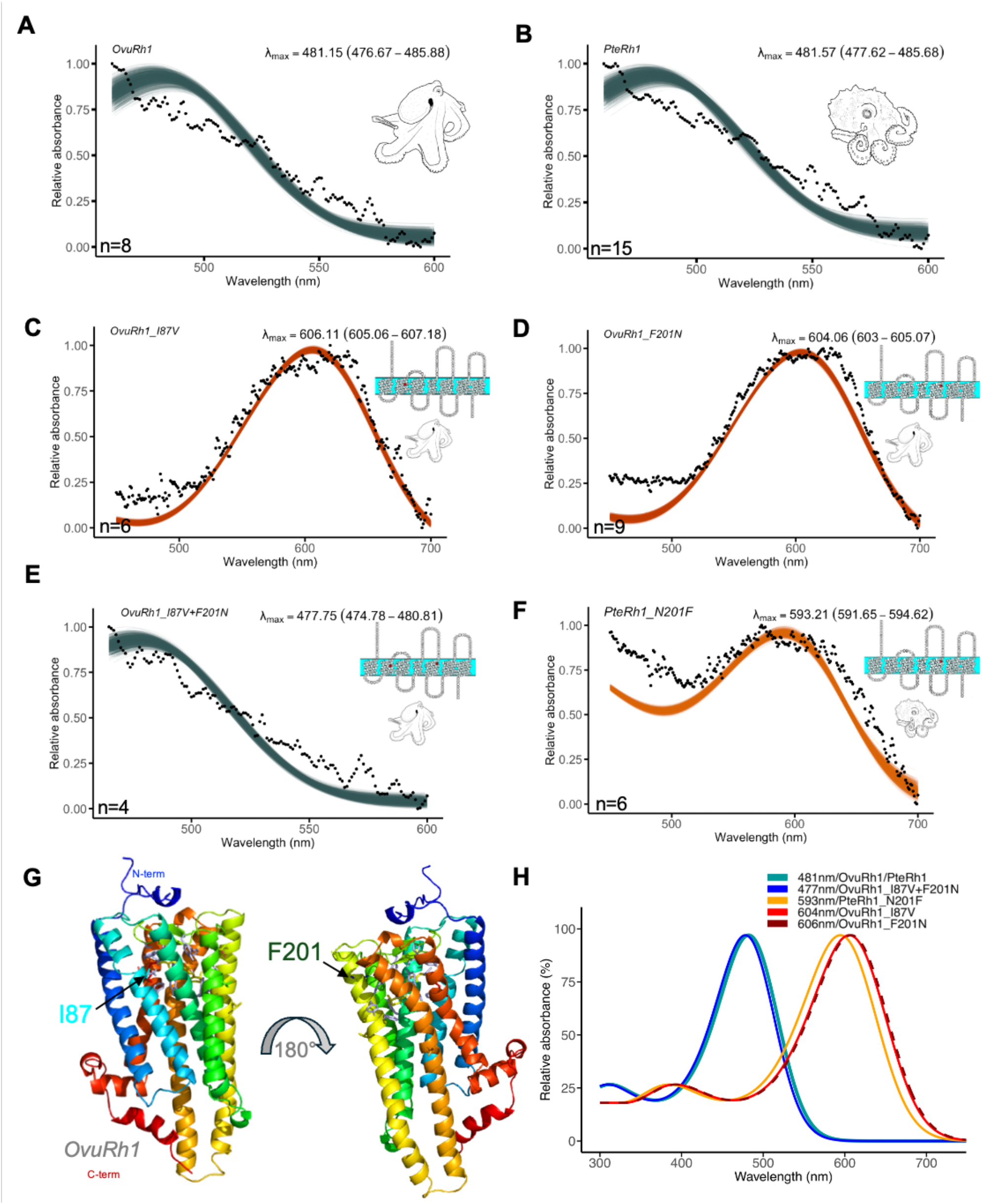
Octopods’ r-opsin1 function as blue-green absorbing visual pigments in littoral and deep-sea species. (A, B) Dark absorption spectrum of *Octopus vulgaris* OvuRh1 and *Pteroctopus tetracirrhus* PteRh1, respectively. The black dots represent the average absorbance across replicates at every wavelength, whereas the lines represent the curve-fitting of individual bootstrap replicates. n is the number of measurements of protein aliquots with active rhodopsin complexes. Absorption spectra of mutated *O. vulgaris* of r-opsin1, with single mutations OvuRh1_I87V (C), OvuRh1_F201N (D), double mutant OvuRh1_I87V+F201N (E) and single mutation of *P. tetracirrhus* r-opsin1 PteRh1_N201F (F). While single mutations show a long-wave shift, the double mutant restores the wild-type Rh1 phenotype. (G) Structure of OvuRh1 with target residues at positions 87 (Helix 2) and 201 (Helix 5). Helices are coloured from blue (N-terminal) to red (C-terminal). (H) Fitted visual pigment functions comparing spectral sensitivities of native and mutant octopus Rh1 opsins. For each rhodopsin, relative absorbance data are fitted to a visual template with polynomial function analyses computed in R to obtain the estimates of lambda max following 1,000 bootstrap replicates. Values in parentheses represent the lower and upper bounds of the 95% confidence intervals. See also accompanying Figures S2-3, File S5.

We then investigated a potential functional role for the substituted residues at positions 87 and 201 between OvuRh1 and PteRh1 (Figure 4C-H, Figure S2B-C). Introducing V87 in the *O. vulgaris* r-opsin1 resulted in a pronounced bathochromic shift of the OvuRh1_V87 dark spectrum with λ_max_ = 606 nm compared with the wild-type OvuRh1 protein (Figure 4C). Similarly, substituting F201 for N201 resulted in a long-wave shifted rhodopsin with λ_max_ = 604 nm (Figure 4D, Figure S2B-C). Both mutant proteins OvuRh1_V87 and OvuRh1_N201 formed protonated, blue-shifted photoproducts (λ_max_ = 429-431 nm) upon acid denaturation (Figure S3A-B), confirming that the red-shifted mutant rhodopsin in the dark state formed a covalent bond to 11-*cis*-retinal. When introducing I87 in PteRh1, no active complexes could be observed, but when substituting F201, PteRh1-N201F absorbed maximally at 593 ± 1.5 nm (Figure 4F), causing a sensibly similar bathochromic shift as OvuRh1-F201N albeit with a broader long-wavelength shoulder of absorbance than observed for the reciprocal F201N mutation in OvuRh1 (Figure 4D). Again, the corresponding PteRh1-N201F rhodopsin acid photoproduct absorbed with a blue-shifted maximal sensitivity at 431 nm (Figure S3C). Interestingly, when both PteRh1 residues are introduced simultaneously in the OvuRh1 backbone, the resulting OvuRh1_V87+N201 dark spectrum absorbed wavelengths at 478 ± 3 nm, within the range of the wild-type rhodopsin of both species (Figure 4E), therefore restoring blue-green absorbance (Figure 4H).

### Ancestral octopods’ r-opsin sensitivity states are slightly green-shifted

We reconstructed r-opsin1 inferred ancestral states across octopod lineages and tested the absorbance spectrum at two key evolutionary nodes: the common ancestor between *Octopus* and *Pteroctopus* (node 9, N9) and the most recent divergent leading to *Pteroctopus* (node 11, N11). These represent the most recent common ancestor to *P. tetracirrhus* and *O. vulgaris*, and the most proximal ancestor to *O. vulgaris*, respectively. The ancestral r-opsin1 at N9 shares 93.3% with PteRh1 (32 distinct residues) and 97% with OvuRh1 (14 distinct residues), whereas the ancestral rhodopsin at N11 shares 95% to both Ovu and Pte Rh1 protein sequences (22 or 23 aa residues differences). Both ancestral node r-opsins1 also bear I87 and F201, and the ancestral residues also corresponding to the OvuRh1 state at otherwise selected sites in PteRh1 (File S1). When reconstituted and purified *in vitro*, these ancestral rhodopsins absorbed in the blue-green spectrum, and maximally at 489 ± 3.3 nm and 492 ± 5 nm, respectively (Figure 5A-D, Figure S2D).

**Figure 5.**
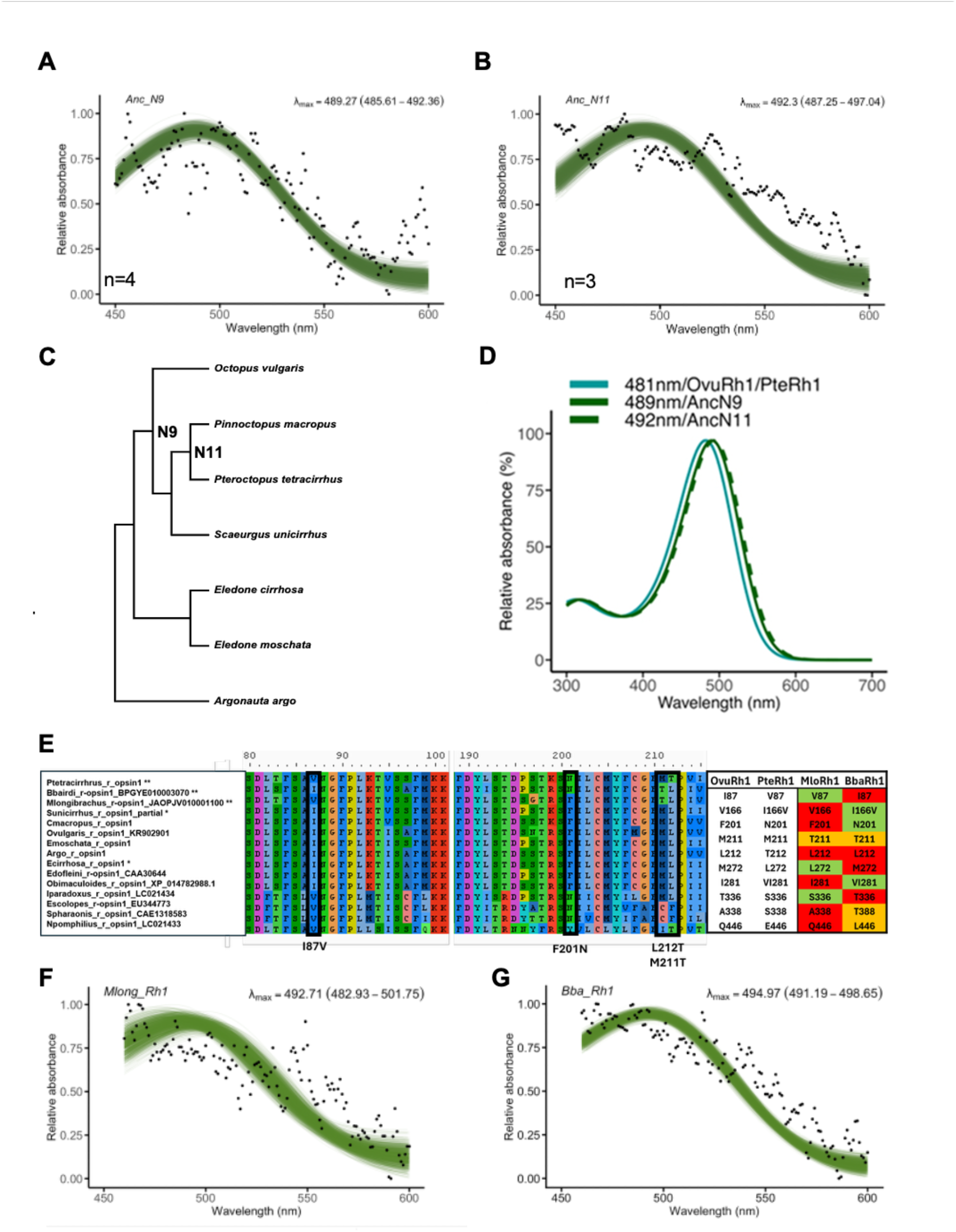
Ancestral octopod rhodopsin absorbance and evolution at homologous PteRh1 sites under positive selection in other species of octopus. sites under positive selection in other species of octopus. (A-B) Dark absorbance spectra for r-opsin1 states at Node 9 (Anc_N9) and Node 11 (Anc_N11), (C) Phylogeny showing the ancestral nodes under investigation, and (D) Comparison between O. vulgaris and *P. tetracirrhus* r-opsin1 absorption spectra and those of the ancestors. (E) The alignment shows the regions showing sites inferred to be under positive selection (BEB > 0.9, highlighted by the black rectangles) of multiple r-opsin1 sequences from different cephalopod species. Site S111 is also indicated due to its proximity to L212T. The table summarizes mutations shared among PteRh1, MloRh1, and BbaRh1. (F-G) Dark absorbance spectra for r-opsin1 states of *M. longibrachus* and *B. bairdii* (n=5). The black dots represent the average absorbance across replicates at every wavelength, whereas the lines represent the curve-fitting of individual bootstrap replicates. The “n” represents the number of measurements of protein aliquots with active rhodopsin complexes (File S5). The * indicates species living from 200 to 500 m, the ** indicates species living more than 500 m of depth. *Architeuthis dux* sequence from De Vivo et al. (2023).

### Atlantic and pacific deep-sea dwelling octopods share homologous substitutions to PteRh1

We subsequently analysed the recently released genomes of two deep-sea octopod dwellers, the Baird’s octopus *Bathypolypus bairdii* and the inkless octopus *Muusoctopus longibrachus* to assess whether r-opsin1 residues under positive selection in *P. tetracirrhus*, including sites 87 and 201, have evolved independently in Atlantic and Pacific lineages (Figure 5E). The former inhabits the Northwest Atlantic Ocean at depths up to 550 m, while the latter is found in the Southeastern Pacific Ocean at depths reaching 1000 m (Jereb et al., 2005). The comparative analysis of their rhabdomeric opsins with other available deep-sea species revealed that, despite the phylogenetic distances, most substitutions found in PteRh1 are present in one or the other species, that is, 8 out of 12 sites (aa 87, 166, 201, 272, 281, 336, 338 and 452) (Figure 5E). In particular, V87 is shared with *M. longibrachus*, whereas N201 is shared with *B. bairdii*. The combination of 87V with 201N is also observed in Decapodiformes, whereas 87V occurs in the r-opsin1 of the chambered nautilus *Nautilus pompilius* (Figure 5E). Intrigued by this intermediate convergent pattern at multiple sites under positive selection, we purified these two additional r-opsin1 and our functional analysis show that these two deep-sea species have conserved broad blue-green rhodopsins compared to those of *O. vulgaris* and *P. tetracirrhus*, with a slightly green-shifted maximal absorption in the 490 nm range (Figure 5F-G).

### Mutation in site 201 decreases opsin adiabatic compressibility in deep-sea species

Hydrostatic pressure can affect protein function, and a reduction in protein compressibility has been proposed as an adaptation to deep-sea environments (Porter et al., 2016). To test whether the mutations identified under positive selection influence protein adiabatic compressibility (βs) of OvuRh1 and PteRh1, we implemented the method from Porter et al. (2016). PteRh1 exhibited lower predicted compressibility (βs = 11.494) than OvuRh1 (βs = 12.005), and therefore a possible adaptation to the deep-sea environment. Our analysis further revealed that the F201N substitution is the major contributor to this reduction in compressibility. Reverting this mutation in PteRh1 to the residue present in all other non-deep-sea species, increased βs to 11.819 (+0.325), indicating that F201N accounts for more than half of the total compressibility difference between PteRh1 and OvuRh1. In contrast, reverting I87 for V87 as in non-deep-sea species contributed a negligible change in βs (11.406 (−0.088)). This effect is consistent with the marked reduction in thermodynamic transfer hydrophobicity (Ht) associated with the substitution of phenylalanine (Phe) by asparagine (Asn) at position 201 (Ht from 2.87 to 0.09), which is predicted to make the region surrounding the retinal-binding pocket less compressible and, therefore, more resistant to hydrostatic pressure. A similarly low compressibility was predicted for BbaRh1 (βs = 11.351). Additional support for the role of residue 201 comes from other deep-sea octopods. Several species of *Bathypolipus* and *Benthoctopus* r-opsin1 possess the N201 substitution and exhibit reduced predicted compressibility (Porter et al., 2016). A similar pattern is observed in the *Thaumeledone* genus r-opsin1 that carry a serine 201 (S), which substantially decreases hydrophobicity, suggesting convergent molecular evolution toward reduced protein compressibility in response to hydrostatic pressure. Although deep-sea Decapodiformes r opsins analysed earlier in Porter et al. (2016) possess both the 87V and 201N substitutions similarly to PterRh1, these specific residues were not predicted to significantly contribute to reduced compressibility in that lineage. This suggests that the structural effects of these substitutions depend on the broader sequence context and may be therefore lineage specific.

## Discussion

In many deep-sea animals, including cephalopods, relatively larger eyes are a common adaptation to low-light environments (Hanke and Kelber, 2020; Warrant and Locket, 2004). The eyes of *P. tetracirrhus,* a mesobenthic octopus living down to depths of 700 m, are notably larger than those of epibenthic species, reaching up to 3 cm in diameter, compared to approximately 2 cm in *S. unicirrhus* and *O. vulgaris*, the latter roughly corresponding to the size of a human eye (Hanke and Kelber, 2020). Given that eyes are energetically costly, this anatomic enlargement likely reflects an adaptation to enhance vision in low-light environments. Deep-sea octopods such as *M. longibrachus* and *B. bairdii* share ecological habitat preferences with *P. tetracirrhus*, and enlarged or prominent eyes (Luna et al., 2021), a morphological adaptation that align with maximizing photon capture in sunlight-depleted habitats, or reliance on other light sources such as prey bioluminescence in low-light environments. This strong energetic investment highlights the importance of studying the evolution of the visual system in these animals, especially in association with extreme environments. Comparing retinal and optic lobe transcripts revealed the prevalence of r-opsin1 as main photoreceptive pigment in the octopod visual system, alongside an overall opsin-expression pattern consistent with the cuttlefish *Sepia officinalis* (Bonadè et al., 2020), the pygmy squid *Idiosepius paradoxus,* and the chambered nautilus *N. pompilius* (Yoshida et al., 2015). Interestingly, among modern cephalopods, peropsin, which has been lost in squids, and cuttlefish (Decapodiformes) appears expressed in several octopods (Figure S1), although its role in the visual system remains to be deciphered. Therefore, cephalopods likely mainly rely on retinal photoreceptors expressing only r-opsin1 and its associated signalling cascade for image-forming vision (Zhang et al., 2021), which together with behavioural tests supports monochromacy as the likely visual model in these animals (Marshall and Cronin, 2011; Marshall and Messenger, 1996; Messenger et al., 1973; Stubbs and Stubbs, 2016).

Cephalopods r-opsin1 genes have not been extensively characterized in heterologous expression systems to validate potential key tuning sites, and our study demonstrates the potential of combining evolutionary, modelling and experimental approaches in this clade. In particular, we measured the absorption spectra of *P. tetracirrhus* and *O. vulgaris* r-opsin1 to test if key amino acid differences may be associated with differences in wavelength absorption. Our results show conservation in r-opsin1 absorption spectra of these deep-sea and littoral species, while single mutations perturb spectral sensitivity, suggesting that the variant sites under selection in *P. tetracirrhus* are associated with multiple functions, leading to compensatory effects that may prevent spectral tuning. In contrast, a 10 nm shift toward longer wavelengths is observed in other analysed deep-sea species, such as *B. bairdii* and *M. longibrachus,* or ancestral molecules (Anc_N9 and Anc_N11). This is an interesting observation that may be indicative of local adaptations, e.g. to perceive variable bioluminescence patterns when increased visual sensitivity to intermitted, low-intensity green light may be advantageous for benthic predators like octopods with limited vertical migration, allowing them to detect prey such as benthic decapods or their hosts, including corals (Johnsen et al., 2012).

Notably, the evolutionary conserved broad rhodopsin spectral sensitivity between ∼470-490 nm is consistent with a proposed secondary epipelagic migration scenario, in which a deep-sea mesopelagic octopod ancestor that gave rise to littoral species (Klug et al., 2023). Epipelagic species are exposed to the full range of visible wavelengths and may therefore evolve shifts in spectral sensitivity to optimize photon capture under different light conditions. In contrast, deep-sea species are restricted to the narrower blue-green range of wavelengths that penetrate or are produced in their environment. Consequently, transitions from shallow to deep habitats are expected to show shifts in spectral sensitivity towards shorter wavelengths, whereas the reverse transition should be less constrained as the blue-green wavelengths available in the deep sea are also present in surface waters. Accordingly, if extant littoral octopods descended from deep-sea ancestor species adapted to dysphotic environments, their ancestral blue-green spectral tuning would already be capable of capturing incoming sunlight when colonizing littoral habitats, without requiring major spectral shifts. The deep-sea evolutionary origin therefore provides a plausible explanation for the remarkably low variation in r-opsin1 spectral sensitivities observed across octopods in our study, in line with earlier electrophysiological measurements (Chung and Marshall, 2016). A second factor may further favour the conservation of the ancestral spectral sensitivity. Independent of evolutionary history, the optical properties of seawater allow wavelengths of light that penetrate water most effectively to propagate over longer horizontal distances, enhancing the detection of distant objects without necessarily imposing strong selective pressure for spectral shifts, even at depths from 500 m and beyond. This constraint is particularly relevant for monochromatic organisms such as those of octopods, which rely heavily on polarization vision rather than colour discrimination (Temple et al., 2021). Because polarization cues are abundant within the blue-green wavelengths dominating underwater light, spectral sensitivity in this range may maximize both photon capture and polarization detection. Consistent with this, littoral species such as *O. vulgaris*, despite being exposed to a broader spectrum of light than deep-sea species may maintain a peak of rhodopsin sensitivity around 475–480 nm possibly reflecting optimization for both photon capture and the detection of polarization cues within the narrower blue-green wavelength band that propagates most efficiently through both horizontal and vertical water layers (Figure 4) (Chung and Marshall, 2016; Inoue et al., 2007).

In addition, the deepest-living octopod species occupying mesobenthic habitats, or those migrating into sunlight-depleted areas of the water column encounter a unique combination of simultaneous ecological constraints, including a reduced photon availability for visual navigation, increasing hydrostatic pressure and intermittent bioluminescence in otherwise near-complete absence of sunlight. Adaptation to high hydrostatic pressures can occur through extrinsic adaptations including amino acid substitutions to cope with cellular and membrane modifications, and intrinsic mechanisms that can reduce protein compressibility (Somero, 1992; Porter et al., 2016). In *P. tetracirrhus*, site 201 considerably decreases PteRh1’s hydrophobicity, and therefore, the adiabatic compressibility in the retinal binding pocket region, supporting its potential contribution as an adaptation to high hydrostatic pressure. Although this residue does not correspond to pressure-related mutations previously identified in the outer regions of cephalopod rhodopsins (Porter et al., 2016), it suggests that alternative molecular mechanisms may underlie pressure adaptation in other species. Together these findings indicate that deep-sea selective pressures may favour amino acid substitutions that preserve rhodopsin stability and function without substantially altering spectral sensitivity. Hence, experimentally, the two validated substitutions (sites 87 and 201), positioned on opposite sides of the retinal binding pocket, contribute to maintaining stable retinal interactions while preserving the ancestral blue-green rhodopsin spectral profile. These selective pressures may explain why, among all Gulf of Naples species, only *P. tetracirrhus,* the deepest-dwelling species in the Gulf, exhibits evidence of positive selection in r-opsin1.

Substituting either PteRh1 N201 to F201 or I87 to V87, as well as one reverse mutation (F201N in OvuRh1), caused unexpectedly large bathochromatic spectral shifts albeit of different magnitude between backgrounds whereas the double mutant in OvuRh1 restored rhodopsin absorbance as in the wild-type PteRh1, supporting that these sites may stabilize the retinal binding pocket, possibly compensating for perturbations in local interactions affecting the chromophore environment. Since those two mutations are at a distance higher than 5Å of the chromophore, and not directly interacting with it, the spectral effect is likely mediated through the surrounding network of conserved residues that shape the chromophore environment. These findings are consistent with previous structural and evolutionary studies implicating homologous rhodopsin residues 87, 88, and 201 in stabilizing the Schiff base, retinal release kinetics and spectral tuning (Chi et al., 2025; Bosh et al., 2003; Sekharan et al., 2012; Varma et al., 2019; Church et al., 2021), supporting the functional relevance of potential compensatory interactions. Both mutations in sites 87 and 201 likely evolved along the branch leading to *P. tetracirrhus* as they are absent in the two phylogenetically related octopod species, *P. macropus* and *S. unicirrhus*. Mutations I87V or F201N also occur naturally in either rhodopsin of other deep-sea octopods, *e.g.* MloRh1 and BbaRh1, respectively, without necessarily causing a large bathochromic shift, but showing a modest ∼ 10 nm shift toward longer wavelengths. This suggests compensatory interactions by other sites stabilising the binding pocket to maintain the rhodopsin blue-green absorbance range. The independent occurrence of these mutations in natural populations is particularly important, as it allows to study their effects in naturally evolved molecules under selective pressures, rather than in artificially engineered constructs, highlighting how the same mutation can produce different outcomes depending on the molecular background. The shared presence of these mutations in deep sea octopods might potentially reflect specific adaptations to low-light environments, including sunlight scarcity and sensitivity to low-intensity intermittent bioluminescence. In addition, as predicted bistable opsins, r-opsins must efficiently revert retinal to the 11-*cis* configuration even under limited light, and some of the mutations observed in deep-sea species may contribute to this distinct process, ensuring reliable photoreception in extreme environments. It is interesting that lineage-specific substitutions were required during the evolution of *P. tetracirrhus* r-opsin1. If F201N evolved primarily as an adaptation to higher hydrostatic pressures of the deep sea by reducing protein compressibility, this substitution may have altered the spectral properties of r-opsin1, necessitating compensatory mutations to restore a broad blue-green sensitivity. Under this scenario, the adaptation of *P. tetracirrhus* r-opsin1 may have involved two interacting evolutionary processes: first, a pleiotropic mutation (F201N) that improved protein performance under high pressure while perturbing spectral sensitivity, and second, compensatory epistasis through I87V, which preserved the adaptive benefit of reduced compressibility while restoring optimal visual pigment function.

In the jumping spider rhodopsin, the negative electrostatic interaction at site Y126 (homologous to Y112 in PteRh1) has been proposed to stabilize the positively charged Schiff base in the 11-*cis* configuration, together with M99 and M103 (homologous to F84 and M88 in PteRh1, respectively, which interact with site 87). The latter two residues are also hypothesized to contribute to Schiff base stabilization when the retinal is in the all-*trans* form (Church et al., 2021; Varma et al., 2019). PteRh1 residues 189 and 205 possibly interact with site 201, these sites are known in cephalopod for their potential role in spectral tuning (reported as sites 202 and 206 in Chung and Marshall, 2016). Furthermore, in squid and vertebrate melanopsins the homologous of PteRh1 site 88, which interact with site 87, contribute to the maintenance of the internal water-mediated hydrogen-bond network required for Schiff base stability, structural cohesion, and efficient photoactivation (Sekharan et al., 2012). The functional importance of site 88 is further highlighted by the fact that its homologous site in human rhodopsin has been linked to the development of retinitis pigmentosa (Bosh et al., 2003). In the branch leading to apes, the rhodopsin site homologous to PteRh1 site 87 has been shown to be under positive selection (F88L) and it is reported to be critical for rhodopsin retinal release rates (Chi et al., 2025), showing that similar molecular sites can undergo parallel evolutionary processes even after hundreds of millions of years of divergence, and showing that this site is important for opsin-retinal interactions.

By integrating transcriptomics, evolutionary analyses, structural modelling, and functional assays, our study shows how r-opsin1 has evolved across octopods living in distinct marine light habitats, underscoring the benefits of combining evolutionary, structural, and experimental approaches to understand how vision has evolved in extreme environments. Despite relying on a single visual opsin that conserves blue-green sensitivity across the group, bathypelagic octopods also exhibit lineage-specific molecular adaptations. In *P. tetracirrhus*, positively selected sites stabilize retinal interactions through molecular context-dependent local epistasis that preserves overall function, and homologous mutations are independently observed in other deep-sea species such as *B. bairdii* and *M. longibrachus*, while maintaining a broad absorption within the blue-green range observed in r-opsin1 for other octopods, suggesting convergent evolution in response to similar ecological pressures as an evolutionary compromise between surface vision and photon capture in low-light, but more bioluminescence-rich habitats. Additionally, residues reflecting adaptation to high hydrostatic pressures emphasize the multifactorial nature of deep-sea adaptation.

Our findings illustrate that evolutionary innovation in a constrained single-opsin visual system such as deep-sea octopods is not achieved through drastic or repeated spectral diversification. Instead, our work illustrates that ecological diversification can be achieved through molecular evolutionary trade-offs that finely tune structural and functional performance to preserve a broadly efficient visual pigment for photoreception in deep-sea habitats.

## Materials and Methods

### RNA-seq of retina and optic lobe

Fresh biological samples were obtained from the octopod species *Eledone moschata*, *Eledone cirrhosa*, *Pinnoctopus macropus*, *Pteroctopus tetracirrhus*, and *Scaeurgus unicirrhus*, collected from fishermen bycatch in the Gulf of Naples. The right eye and optic lobe were dissected from each specimen, and tissues were preserved in RNAlater (Sigma) at -20°C prior to total RNA extraction. Following cDNA library preparation, transcriptomic analyses were performed using Ion Torrent next-generation sequencing (NGS) technology. Raw sequencing reads were quality-checked with FastQC v0.12 and trimmed using Trimmomatic v0.38 (parameters: LEADING:5 TRAILING:5 SLIDINGWINDOW:4:15 MINLEN:50) to remove low-quality sequences and bases (Table S1). *De novo* assembly was performed using Trinity v2.15 (parameters: --min_kmer_cov 2 --normalize_reads; Haas et al., 2013) (Table S2).

### Opsin search, cloning and expression analysis

Opsin genes were retrieved from the assembled transcriptomes using BLASTn, with *Octopus vulgaris* opsin sequences as query. Their identity was confirmed by reverse BLASTn and by visual inspection of their sequence alignment. A similar approach was applied to identify the *Argonauta argo* opsin coding sequences (Yoshida et al., 2022). r-opsin primers were designed to amplify the full-length sequences via reverse-transcription PCR (RT-PCR). The resulting products were cloned into the pGEM®-T Easy Vector, plasmid DNAs were purified and verified through Sanger sequencing.

Gene expression analysis was conducted using the Trinity v.2.15 Transcript Quantification protocol, which supports alignment-based methods such as RSEM (Grabherr et al., 2011; Li and Dewey, 2011). Expression profiles of opsin genes were plotted using ggplot2 R v. 3.6.6 (Wickham, 2009). Data on expression analysis in *Octopus vulgaris* have been obtained by combining the optic lobe and retina transcript short reading assembly (SRR17321848 and SRR17321849) mapped on CDS data (GCA_951406725.2) available on NCBI before quantification.

### *Octopus* phylogeny and evolutionary analyses

The species phylogeny was reconstructed using the collected r-opsin1 sequences with IQ-TREE2 (model: -m MF -B 1000) (Minh et al., 2020). Bootstrap values and branch lengths were removed for simplification. The phylogeny was rooted using *Argonauta argo* and is consistent with previous phylogeny (López-Córdova et al., 2022; Sánchez-Márquez et al., 2022; Taite et al., 2023). To reconstruct ancestral r-opsin1 sequences, we implemented baseML (noisy = 3; verbose = 0; runmode = 0; model = 7; Mgene = 0; clock = 0; fix_kappa = 0; kappa = 2.5; fix_alpha = 1; alpha = 0; Malpha = 0; ncatG = 5; fix_rho = 1; rho = 0.; nparK = 0; nhomo = 0; getSE = 0; RateAncestor = 1; Small_Diff = 1e-6; cleandata = 0; icode = 11; fix_blength = 2; method = 0) in PAML (Álvarez-Carretero et al., 2023) (File S1). To assess the selective pressures acting on the visual r-opsin in octopods inhabiting different light environments (*O. vulgaris*, *E. moschata*, *E. cirrhosa*, *P. macropus*, *S. unicirrhus*, *P. tetracirrhus*, *and A. argo*) we performed positive selection analysis in PAML codeML. We computed first the one-ratio model (M0; noisy = 0; verbose = 0; runmode = 0; seqtype = 1; CodonFreq = 3; model = 0; NSsites = 0; icode = 1; getSE = 0; fix_blength = 2), which assumes a single dN/dS (ω) ratio across all sites and branches to serve as a null hypothesis. Next, to check if selective pressures occurred in littoral, deep-sea or in single species, M0 was compared to the alternative branch models: M1, in which each branch has its own ω value (noisy = 0; verbose = 0; runmode = 0; seqtype = 1; CodonFreq = 3; model = 1; NSsites = 0; icode = 1; getSE = 0; fix_blength = 2); and M2, where one group of branches (e.g. littoral or deep-sea) which is assumed to experience different selective pressure and therefore have a different ω (noisy = 0; verbose = 0; runmode = 0; seqtype = 1; CodonFreq = 3; model = 2; NSsites = 0; icode = 1; getSE = 0; fix_blength = 2).

Subsequently, we repeated the analysis implementing the branch-site models to test for positive selection on specific branches and sites. Specifically, we compared BSM0 (noisy = 0; verbose = 0; runmode = 0; seqtype = 1; CodonFreq = 3; model = 2; NSsites = 2; icode = 1; getSE = 1; fix_blength = 2; fix_omega = 1; omega = 1), that restricts ω ≤ 1 in all sites to different BSM1 (alternative model; noisy = 0; verbose = 0; runmode = 0; seqtype = 1; CodonFreq = 3; model = 2; NSsites = 2; icode = 1; getSE = 1; fix_blength = 2; Small_Diff = 0.1e-6), that allows ω > 1 at specific sites in foreground branches.

The comparison was performed using the likelihood ratio test (LRT). Reported results include the log-likelihood difference (2ΔlnL), degrees of freedom (DoF), and p-values (see Table 1). Subsequently, the ancestral state have been reconstructed under PAML using baseML (noisy=3 verbose=0 runmode=0 model=7 Mgene=0 clock=0 fix_kappa=0 kappa=2.5 fix_alpha=1 alpha=0 Malpha=0 ncatG=5 fix_rho=1 rho=0 nparK=0 nhomo=0 getSE=0 RateAncestor=1 Small_Diff=1e-6 cleandata=0 icode=11 fix_blength=2 method=0).

### Mapping sites under positive selection

To examine whether the sites under positive selection in PteRh1 may be associated with spectral tuning, we mapped them onto a 3D model of the r-opsin1 protein generated using the I-TASSER server On-line Server (Zheng et al., 2025) (File S2) and visualized in ChimeraX (Meng et al., 2023), then measured the distances between selected residues and the retinal chromophore. Site numbering corresponds to the PteRh1 opsin, and r-opsin sequences were also aligned between cephalopods, the jumping spider and the cow rhodopsin using MAFFT to provide corresponding reference numbering (File S2). To double confirm these results, we performed aBSREL analysis (2.6) on the alignment from /home/datamonkey/datamonkey-js-server/app/absrel using HyPhy v2.5.98 (Smith et al., 2015).

### Analysis on adiabatic compressibility of octopod opsin

We implemented the method described by Porter et al. (2016), which is based on the regression model developed by Gromiha and Ponnuswamy (1993), to estimate protein adiabatic compressibility (βs). This sequence-based approach predicts protein compressibility by incorporating the physicochemical properties of amino acids. We calculated βs for both the transmembrane and loop regions (amino acids 32–324), as well as for the full-length protein.

To assess the contribution of individual amino acids, we additionally evaluated the effect of single amino acid substitutions by calculating the change in βs resulting from each substitution. The equations of Gromiha and Ponnuswamy (1993) were implemented in a Microsoft Excel workbook with the assistance of Claude (Anthropic), and all equations and parameter values were subsequently verified by manual inspection to ensure fidelity to the original publication (see Supplementary Information).

### Opsin cloning and construct preparation

Coding sequences for *O. vulgaris* (Ovu), *P. tetracirrhus* (Pte), *B. bairdii* (Bba) and *M. longibrachus* (Mlo) Rh1 genes, and ancestral sequence reconstructions (ASR) were flanked by HindIII and BsiWI restriction sites and a GTCGCC Kozak sequence in 5’ prior codon optimization for mammalian expression in pUC57 plasmids (Genscript). Plasmid DNAs were digested at HindIII/BsiWI (NEB), separated by gel electrophoresis, gel purified using the Monarch DNA gel extraction kit (NEB), and ligated with T4 DNA ligase in the linearized pcDNA5-FLAG-T2A-mRuby2 expression vector (Lienard et al., 2021). Plasmid DNAs were transformed into chemically competent NEB 5-alpha cells (NEB) and single bacterial clones were cultured overnight in Luria-Broth (LB) containing 100 µg/mL Ampicillin, purified using the Monarch Plasmid Miniprep Kit (NEB), and verified by Sanger Sequencing (Table S3). Larger bacterial cultures were prepared for final constructs to prepare stock plasmid DNAs following the Endotoxin-free plasmid midi procedure (Machery-Nagel).

### Homology-modelling and site-targeted mutagenesis

OvuRh1 and PteRh1 rhodopsin homology structures were generated in Swiss-Model (Schwede, 2003) against PDB:2z73 and visualized in PyMol (Rigsby and Parker, 2016). Of the 32 amino acid substitutions between both rhodopsins, 21 conserved sites were predicted to lie within 5Å of the retinal (Table S4). We also mapped the position of other 5 variant amino acids (S1-S5) identified to be under selection relative to their helix position and predicted vicinity with sites lining the binding pocket. To test for potential effect on Rh1 spectral sensitivity, oligonucleotide primers (Table S3) were designed using the NEBbasechanger tool. Plasmid DNA constructs for OvuRh1 and PteRh1 were modified to incorporate site-specific substitutions using the NEB mutagenesis kit following the manufacturers’ instructions followed by transformation, Sanger sequencing verification and plasmid stock preparation as described above.

### Opsin heterologous expression in HEK203T cells

#### Small-Scale Clone Selection, Transfection, and Functional Assays

Total rhodopsin protein expression for Rh1 wildtype or mutant constructs was validated through small-scale transient transfection and expression optimization in human embryonic kidney (HEK293T) cells. Cells were seeded at a density of 0.6 × 10⁶ per well in 6-well culture plates containing Dulbecco’s modified Eagle medium (DMEM) with High Glucose (Biowest) supplemented with 10% fetal bovine serum (FBS), and transfected at ∼80% confluency (after 24 h) using a 1:2 DNA (3 μg): Polyethylenimine (6 μL of 1 mg/mL in water, PEI MAX, 40,000MW, PolySciences) ratio diluted in Opti-MEM Reduced Serum (Life Technologies). On average, 3 to 4 plasmid DNAs were screened by construct. After 48 h, fluorescence of mRuby2 was assessed under an ECHO revolve microscope. The supernatant was discarded, and cells were harvested in 1 mL cold D-PBS (Sigma-Aldrich) then centrifugated at 4,000 rpm for 5 min at 4°C. The cell pellet was lysed in 30 μL cold RIPA buffer (Invitrogen) supplemented with 1% n-Dodecyl β-D-maltoside (Sigma-Aldrich) and EDTA-free protease inhibitors (Sigma). Lysis was performed for 1h at 4°C with gentle rotation, followed by centrifugation at 13,000 rpm for 15 min at 4°C. The resulting supernatant (crude lysate) was quantified using bovine serum albumin (BSA, Sigma-Aldrich).

Aliquots containing 25 μg of total protein were mixed with 4× Laemmli buffer containing 10% β-mercaptoethanol, incubated at room temperature for 20 min, and resolved via SDS-PAGE (4-15% MP TGX Gel, BioRad) at 80 V for 90 min on ice. Proteins were transferred to nitrocellulose membranes using a TurboBlotTransfer system (Bio-Rad), blocked for 1h with 5% milk in Tris Buffered Saline containing 0.1% Tween 20 (TBS-T), and probed overnight at 4°C with anti-FLAG antibody (1:1,000, #F1804, Sigma-Aldrich) under gentle rocking. After 3x 10 min TBS-T washes, membranes were incubated for 1h at RT with HRP-conjugated ECL anti-mouse antibody (1:2,500, #NA931, Amersham), then washed 3x 10 min in TBS-T. Membrane signals were visualised via chemiluminescence using SuperSignal West Femto (Invitrogen) on an ImageQuant 800 (Amersham). Higher-expressing rhodopsin constructs identified in this screen were selected for large-scale *in vitro* purification.

#### Large scale expression and purification

Larger scale expression and purification followed the general PASHE procedure (Liénard et al., 2021; Liénard et al., 2022) with specific optimization. On day 1, thirty 10-cm dishes were seeded at 2.0 × 10^6^ cells with HEK293T cells. Each plate was transfected on day 2 with lipid complexes formed in serum-reduced Opti-MEM (Life Technologies) containing 24 µg Endotoxin-free plasmid DNA and 48 µL PEI. All steps from here on were carried out in a dark room. Six hours post transfection, under dim red-light illumination, 11-*cis*-retinal was added to each plate at a final concentration of 5 µM (in 100% Ethanol) together with fresh culture medium change. Plates were wrapped in aluminium foil. Forty-eight hours after transfection, cells were collected, washed and resuspended in 10 ml cold resuspension buffer containing 3 mM MgCl_2_, 50 mM Hepes (pH 7.0), 140 mM NaCl and EDTA-free protein inhibitors (Sigma-Aldrich), and nutated for 1 h with 40 µM 11-*cis*-retinal under gentle tube inversion (10 rpm). After centrifugation at 4°C for 25 min at 21,500 rpm in a Optima L-90-K centrifuge (rotor Ti-70, Beckman Coulter), cell membranes were solubilized in 10 mL Hepes buffer supplemented with 1% DDM and 25% Glycerol vol/vol for 1h at 4°C. The crude protein extract (supernatant) was collected after a second round of ultracentrifugation and mixed with 500 µL Pierce Anti-DYKDDDDK Affinity Resin (Thermo Scientific) then incubated at 4°C overnight under gentle rotation in a black sterile 15 mL tube. The next day, the resin-bound fraction was loaded on a 5 mL Pierce column, then rinsed 3x in 3 mL cold wash Hepes buffer containing 25% Glycerol vol/vol, 0.1% DDM without protease inhibitors. The Opsin-FLAG bound fraction was eluted in 1000 uL wash Hepes buffer via competitive binding by addition of 0.7 mg FLAG peptide with gentle rotation for 20-30 min. All centrifugation steps took place at 15°C and 1,000 *g* for 1 min. The total eluate fraction was concentrated at 4°C and 4,000 rpm for 70 min using a 2-mL Amicon Ultra-2 filter (10 kDa) column to a final volume of approx. 150 µL and divided in black 1.5 mL sterile tubes on ice. Aliquots of each purification fractions (flow-through, wash, filtrate and eluate) were kept for SDS-page protein electrophoresis and western blot analyses as described above.

#### Absorbance measurements under dim red-light

To obtain dark spectra, UV-Vis wavescans (250-750nm) from 1-2 µL eluate fractions were measured on an ImplenN60 spectrophotometer. The production of a deprotonated photoproduct (M state) was monitored via acid denaturation (Tejero et al., 2024). For this, 7 µL of dark-adapted protein eluate were gently mixed with 3 µL of HCL 133 mM in water in a dark 1.5 mL tube, followed by UV-Vis measurements. For all purifications, the conversion of 11-*cis*-retinal into retinal oxime, was verified by mixing 16 µL of dark-adapted opsin eluate with 0.5 µL Hydroxylamine (Sigma-Aldrich), followed by illumination of the whole sample by bright light for 30 s, then UV-vis measurements at 30-sec intervals. For each rhodopsin state, we estimated λ_max_ by nonlinear fitting of the absorbance data using a visual pigment template (Govardovskii et al., 2000). We performed 1,000 bootstrap replicates to compute λ_max_ estimates and the associated confidence intervals in R v. 3.6.6. using the rsample (Frick et al., 2026) and tidymodels (Kuhn and Wickham, 2020) packages.

## Acknowledgements

We thank the SZN Computational Facilities and the RIMAR-BAC service, in particular Pasquale De Luca and Luca Ambrosino for technical help with transcriptomic analysis, Lorena Buono and Tosca van Gelderen for providing Blast script and advice for transcriptomics analysis, Rafael Zardoya and Roberto Feuda for their insightful suggestions, Valérie Panneels for helpful advice on r-opsin 1 expression, Megan Porter for her valuable advice on protein compressibility analysis, and the National Eye Institute for providing 11-*cis*-retinal. We thank OCEANA for the having offered, free of cost, the use of the digital photo of a *Pteroctopus tetracirrhus* -EUO © OCEANA 58773-, to create the schematic drawing for the manuscript.

## Supplementary Material

All raw data and dataset generated in this study are provided as supplementary files accompanying the paper or are available on https://github.com/Xodroont/Deep_sea_Sel. Sequencing data sets are available in the NCBI BioProject database under accession number PRJNA1495468.

## Supplementary Information for

This file contains:

Supplementary Files S1 to S6 can be found at: https://github.com/Xodroont/Deep_sea_Sel

## Supplementary Figures

**Figure S1.**
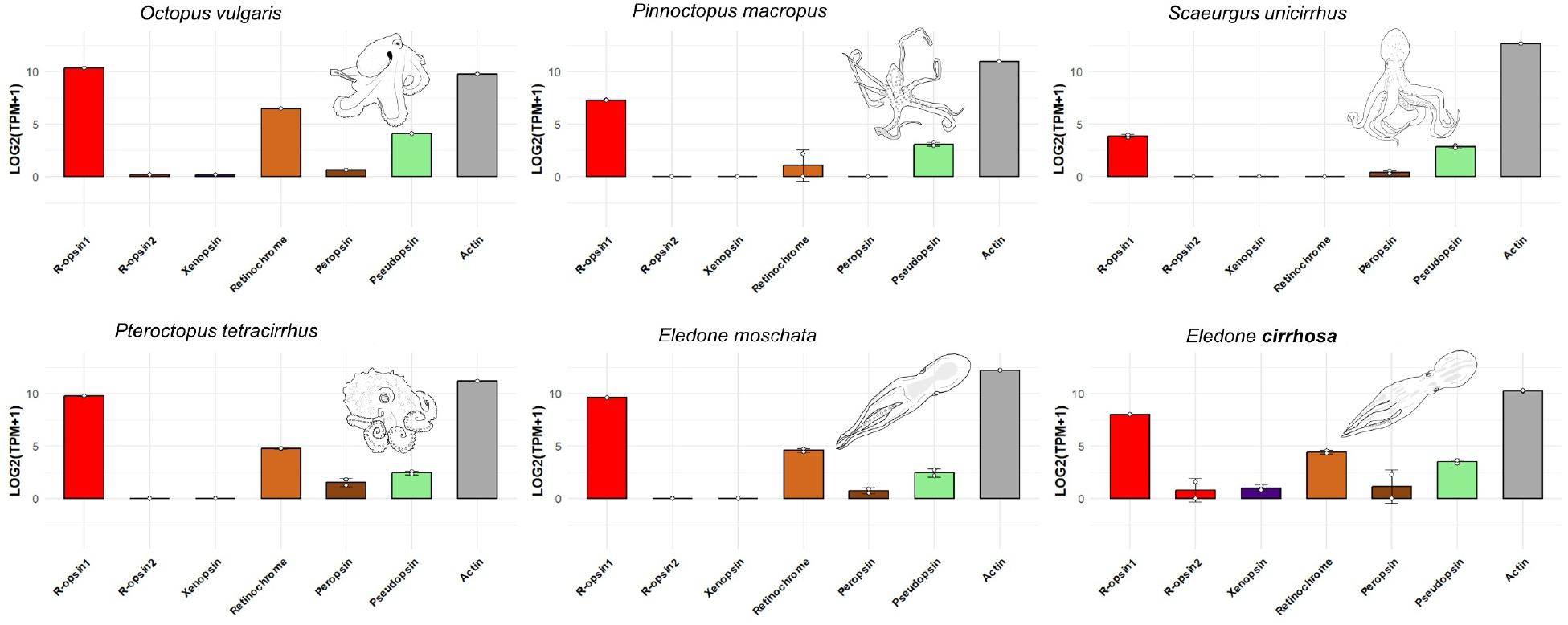
Relative opsin expression in the octopod visual system. Bar plots indicate the presence and relative expression levels of different opsins across six octopod species. r-opsin1 is the most highly expressed opsin in the retina and optic lobe, highlighting its central role in octopod vision. r-opsin1 is the most consistently expressed opsin in the visual system, followed by retinochrome, a cephalopod-specific photoisomerase, that serves as photosensitive receptor with opposite retinal isomerization properties (all-*trans* to 11-*cis*) (Vöcking et al., 2021). Overall, there is generally low expression of xenopsin, r-opsin2, peropsin, and pseudopsin transcripts in the visual system across octopus. See also File S4.

**Figure S2.**
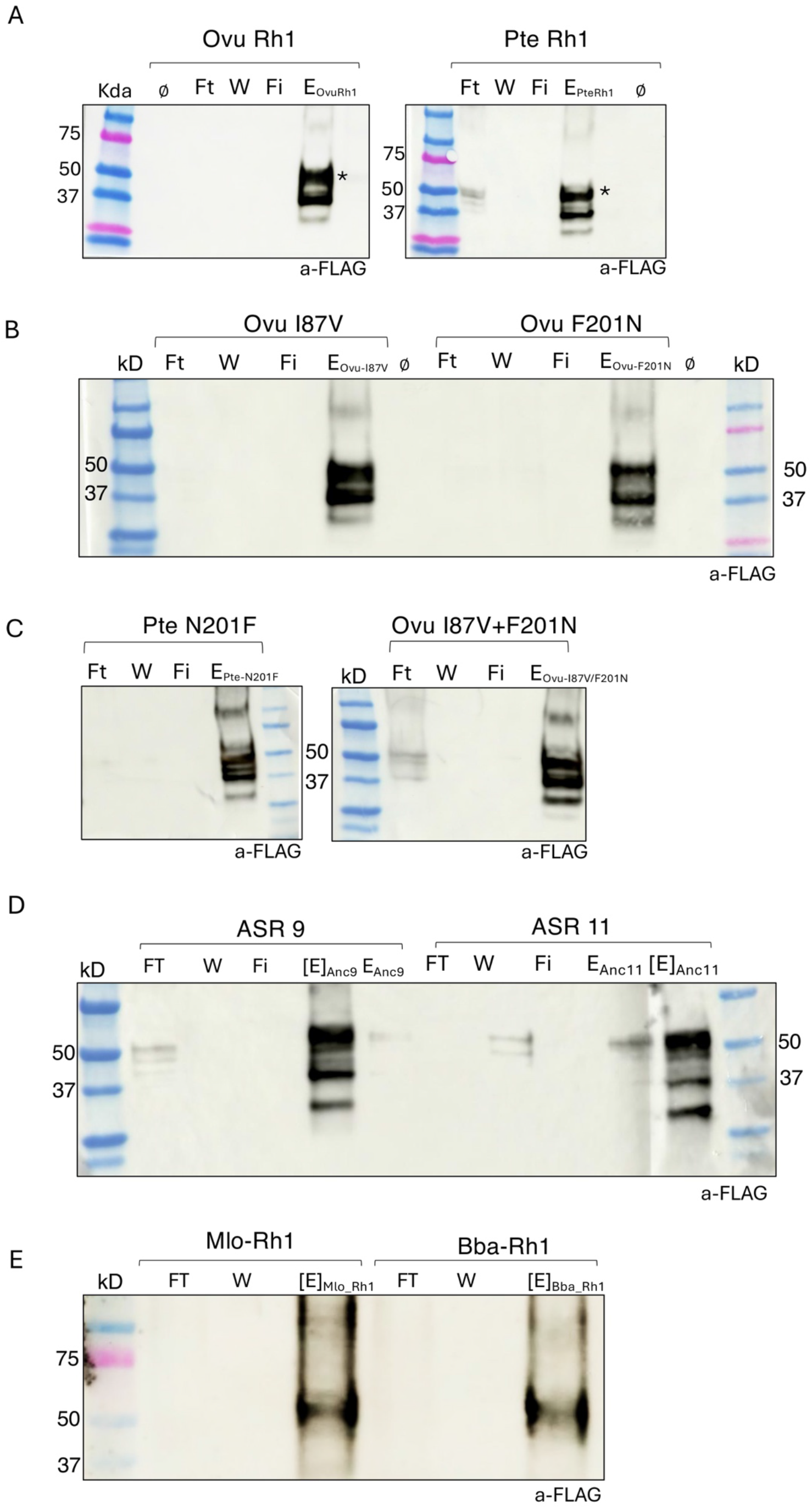
Western Blot analyses opsin purification fractions. FT, Flow through fraction containing non-FLAG opsins, and excess unbound 11-*cis*-retinal. W, wash fraction to remove any residual unbound retinal in solution. Fi, Flow through from the Amico Ultra concentration column. E, non-concentrated eluate. [E], concentrated eluate after centrifugation in the Amicon column. (A) OvuRh1 and PteRh1, asterisks highlight expected monomeric rhodopsin sizes, (B) OvuRh1 I87V (left) and OvuRh1 F201N (right). (C) PteRh1 N201F (left), and OvuRh1 I87V+F201N (right). (D) Ancestral Rh1 sequence reconstruction at nodes 9 (ASR 9) and 11 (ASR 11). (E) Mlo Rh1 and Bba Rh1.

**Figure S3.**
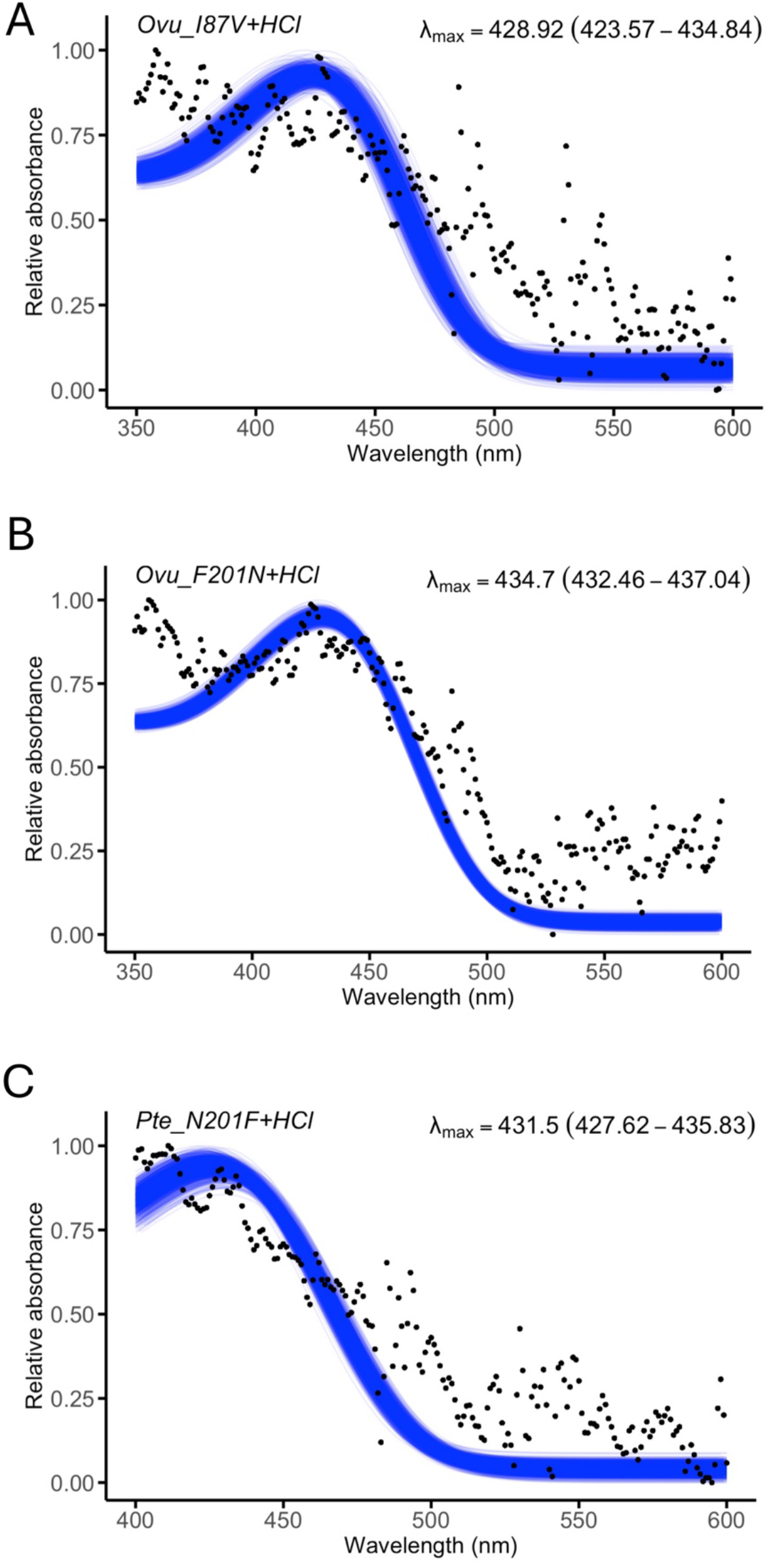
Photoproducts for opsin mutants following acid denaturation. OvuRh1_I87V (A), OvuRh1_F201N (B) and PteRh1_N201F (C). Each purified rhodopsin in complex with 11-*cis*-retinal (Figure 4E-F) was treated with HCl 133 mM and exposed to light (see methods). See accompanying File S5.

## Supplementary Tables

**Table S1.**
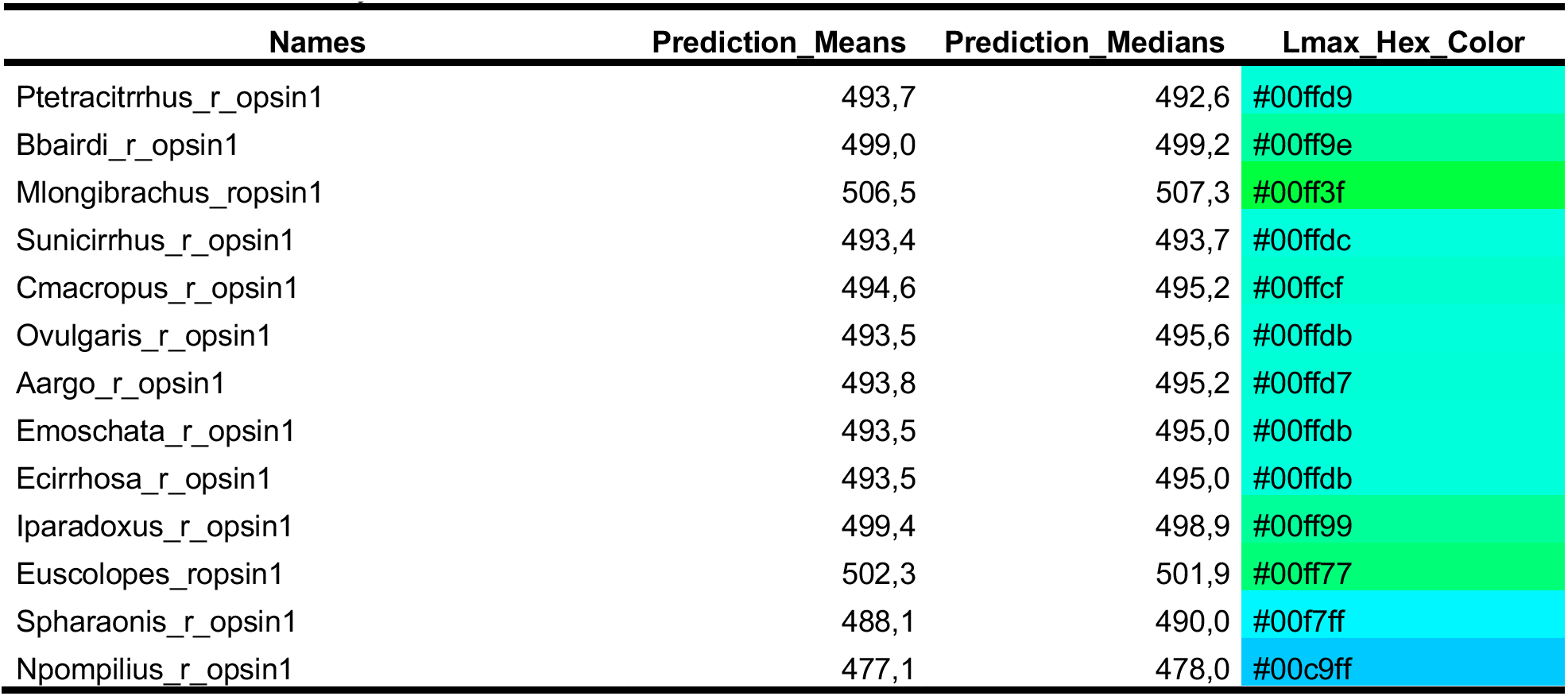
VPOD analysis.

**Table S2.**
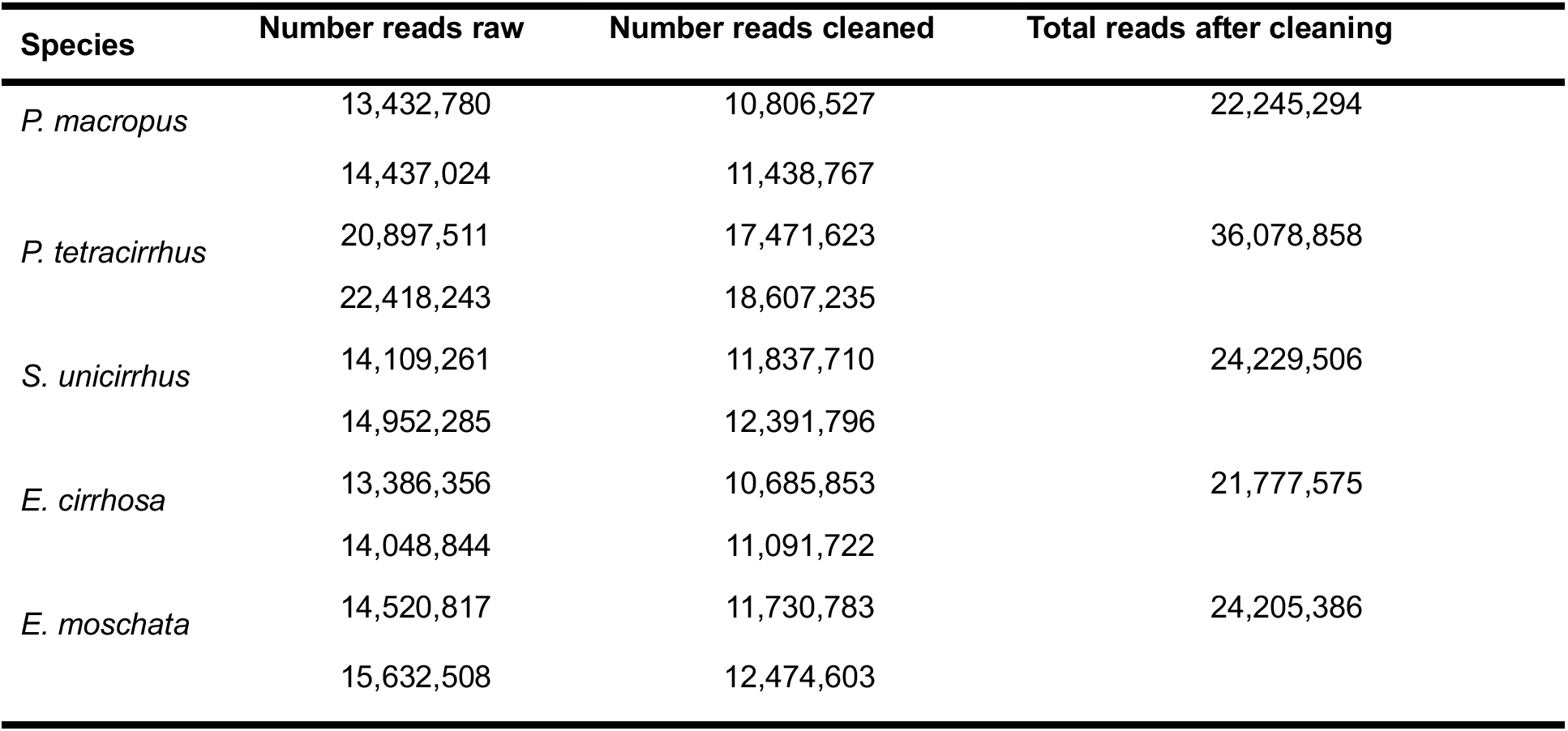
Transcriptomes statistics of five octopod species. The table shows the number of total reads for each species obtained before and after cleaning procedure.

**Table S3.** Prediction of residues in the binding pocket (5A) for *O. vulgaris* and *P. tetracirrhus* Rh1. Those residues are conserved in *B. bairdii* and *M. longibrachus*.

| Residue location | Amino acid position | Amino acid |
| --- | --- | --- |
| TM2 | 84 | Phe (F) |
| TM2 | 88 | Asn (N) |
| TM3 | 112 | Tyr (Y) |
| TM3 | 113 | Gly (G) |
| TM3 | 116 | Gly (G) |
| TM3 | 117 | Gly (G) |
| TM3 | 121 | Phe (F) |
| beta-sheet | 178 | Tyr (Y) |
| beta-sheet | 181 | Glu (E) |
| beta-sheet | 186 | Ser (S) |
| beta-sheet | 187 | Cys (C) |
| beta-sheet | 188 | Ser (S) |
| beta-sheet | 189 | Phe (F) |
| TM5 | 205 | Met (M) |
| TM5 | 206 | Tyr (Y) |
| TM5 | 209 | Gly (G) |
| TM5 | 210 | Phe (F) |
| TM6 | 275 | Trp (W) |
| TM6 | 278 | Tyr (Y) |
| TM6 | 279 | Ala (A) |
| TM7 | 302 | Val (V) |
| TM7 (Bov Rh1 Lys 296) | 306 | Lys (K) |

\* based on original position in *O. vulgaris* and *P. tetracirrhus* Rh1

**Table S4.**
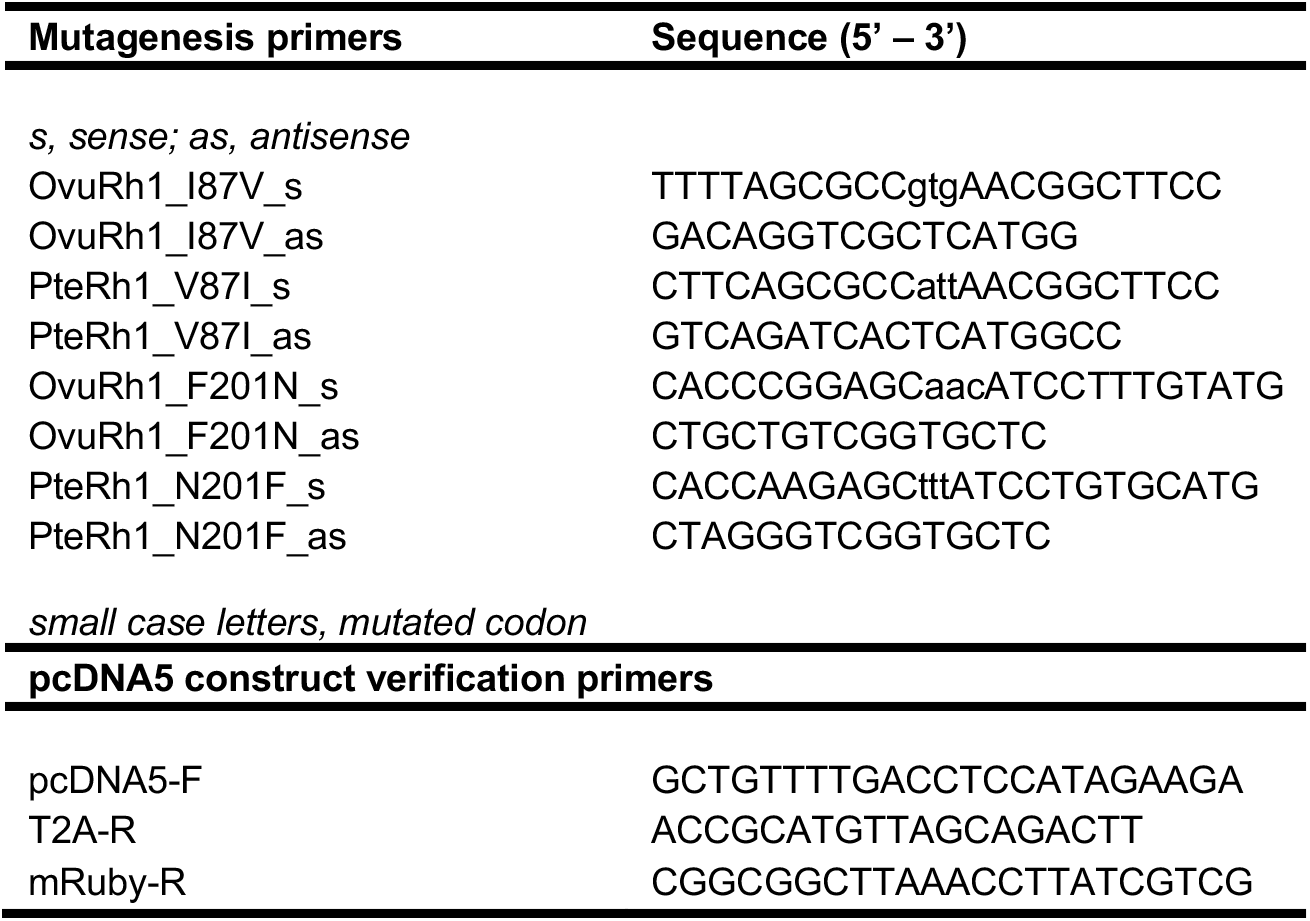
Oligonucleotide sequences used in the present study.

**Adiabatic compressibility (β_s) across octopods r-opsin1**

**Table S5.** Differences in β_s between *P. tetracirrhus* r-opsin and that of littoral octopod species. Two analysis regions are used: (i) full protein and (ii) TM core + loops (residues 32–324). β_s is expressed in units of ×10⁻¹² cm² dyn⁻¹ (equivalent to ×10⁻⁵ MPa⁻¹). A negative Δβ_s indicates that *P. tetracirrhus* is less compressible (stiffer) than the littoral species.

| Sequence | N aa | $\beta_s$ Full protein | $\beta_s$ TM core |
| --- | --- | --- | --- |
| *O. vulgaris* (littoral) | 455 | 3.8473 | 12.0054 |
| *P. macropus* (littoral-control) | 457 | 3.8394 | 11.9578 |
| *P. tetracirrhus* (deep sea) | 455 | 3.2075 | 11.4939 |
| $\Delta\beta_s$ (Pt – avg Lit) | | -0.636 | -0.488 |

**Sequences:**

>P_tetracirrhus_r-opsin1

MVESTTLVNQTWWYNPTVYIHPHWAKFDPIPDAVYYSVGIFIGVVGIIGILGNGLVIFLFSKTKSLQTPANMFIINLAMSDLTFSAVNGFPLKTVSSFMKKWIFGKVACQLYGLLGGIFGFMSINTMAMISIDRYNVIGRPMAASKKMSHRRAFLMIIFVWMWSIIWAVGPVFNWGAYVPEGILTSCSFDYLSTDPSTKSNILCMYFCGFMTPVIIIGFCYFNIVMSVSNHEKEMAAMAKRLNAKELRKAQAGQSAEMKLAKISMVIISQFLLSWSPYAIVALLAQFGPAEWVTPYAAELPVLFAKASAIHNPIVYSVSHPKFREAIQVKFPWLLSCCQFNEKECEDANEAEEEVPASEGGGGESADAAQMKEMMAMMQKMQAQQSAYPQPPPQGYPPQGYPPQGAYPPQGYPPQGAYPPQGYPPQGYPPQGAPPQGEPTQGAPPEGVDNQAYQA

>P_macropus_r-opsin1

MVESTTLVNQTWWYNPTVDIHPHWAKFDPIPDAVYYSVGIFIGVVGIIGILGNGLVIYLFSKTKSLQTPANMFIINLAMSDLSFSAINGFPLKTVSSFMKKWIFGKVACQLYGLLGGIFGFMSINTMAMISIDRYNVIGRPMAASKKMSHRRAFLMIIFVWMWSIVWAVGPVFNWGAYVPEGILTSCSFDYLSTDSSTKSFILCMYFCGFMLPVVIIAFCYFNIVMSVSNHEKEMAAMAKRLNAKELRKAQAGQSAEMKLAKISMVIITQFMLSWSPYAIIALLAQFGPAEWVTPYAAELPVLFAKASAIHNPIVYSVSHPKFREAIQVKFPWLLTCCQFNEKECEDANDAEQEVPASEGGGGGGESADAAQMKEMMAMMQKMQAQQAAYPQPPPQGYPPQGYPPQGAYPPQGYPPQGAYPPQGYPPQGYPPQGAPPQGEATQGAPPQGVDNQAYQA

>O_vulgaris_r-opsin1

MVESSTLVNQTWWYNPTVDIHPHWAKFDPIPDAVYYSVGIFIGVVGIIGIFGNGVVIYLFSKTKSLQTPANMFIINLAMSDLSFSAINGFPLKTISAFMKKWIFGKVACQLYGLLGGIFGFMSINTMAMISIDRYNVIGRPMAASKKMSHRRAFLMIIFVWIWSIVWSVGPVFNWGAYVPEGILTSCSFDYLSTDSSTRSFILCMYFCGFMLPIIIIAFCYFNIVMSVSNHEKEMAAMAKRLNAKELRKAQAGQSAEMKLAKISMVIITQFMLSWSPYAIIALLAQFGPAEWVTPYAAELPVLFAKASAIHNPIVYSVSHPKFREAIQNTFPWLLTCCQFNEKECEDANEAEQEVAPSEGGGGESADAAQMKEMMAMMQKMQAQQAAYQQPPPQGYPPQGYPPQGAYPPQGYPPQGAYPPQGYPPQGYPPQGAPPQGDATQGAPPQGVDNQAYQA

**Equation used: equation 5** (Gromiha & Ponnuswamy, 1993)

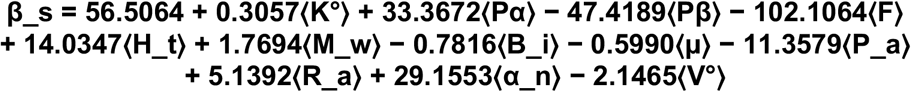

⟨p⟩ = Σ_i f_i · p_i (composition-weighted mean, mole fractions). Equation statistics: R = 0.9546, SE = ±0.8284 ×10⁻¹² cm² dyn⁻¹.

Values are taken from Gromiha & Ponnuswamy (1993) Table 1, with property data sourced from Iqbal & Verrall (1988) Table I. Two properties require unit conversion to match the regression:

- M_w: listed in Da (e.g. Ala = 89) — used as kDa (0.089) in eqn 5
- V°: listed in cm³/mol (e.g. Ala = 60.46) — used as mL/g = V°(cm³/mol) ÷ M_w(g/mol) = 0.679 for Ala

An excel file has been provided as File S6.

**Table S6.**
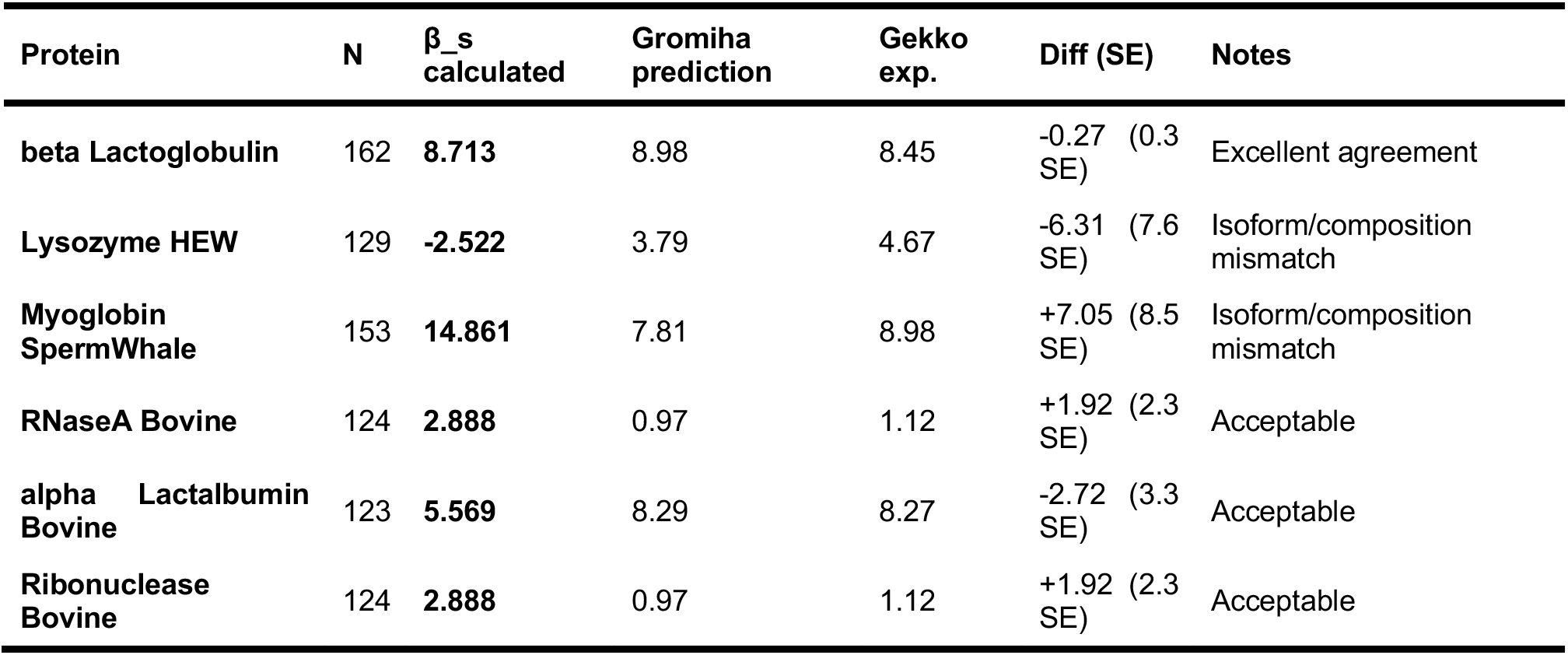
Validation of the β_s equation (Gromiha & Ponnuswamy, 1993, eqn. 5) using β-lactoglobulin (UniProt P02754, mature chain, 162 aa) gives β_s = 8.71 vs Gromiha and Ponnuswamy prediction 8.98 and Gekko & Hasegawa experimental 8.45 (Biochemistry, 1986). Discrepancies observed for other validation proteins reflect differences in amino acid composition between modern database sequences and the historical compositions used by Gromiha and Ponnuswamy (1993).

| Protein | N | $\beta_s$ calculated | Gromiha prediction | Gekko exp. | Diff (SE) | Notes |
| --- | --- | --- | --- | --- | --- | --- |
| beta Lactoglobulin | 162 | 8.713 | 8.98 | 8.45 | -0.27 (0.3 SE) | Excellent agreement |
| Lysozyme HEW | 129 | -2.522 | 3.79 | 4.67 | -6.31 (7.6 SE) | Isoform/composition mismatch |
| Myoglobin SpermWhale | 153 | 14.861 | 7.81 | 8.98 | +7.05 (8.5 SE) | Isoform/composition mismatch |
| RNaseA Bovine | 124 | 2.888 | 0.97 | 1.12 | +1.92 (2.3 SE) | Acceptable |
| alpha Lactalbumin Bovine | 123 | 5.569 | 8.29 | 8.27 | -2.72 (3.3 SE) | Acceptable |
| Ribonuclease Bovine | 124 | 2.888 | 0.97 | 1.12 | +1.92 (2.3 SE) | Acceptable |

**Sequences:**

**beta Lactoglobulin (162 aa):**

LIVTQTMKGLDIQKVAGTWYSLAMAASDISLLDAQSAPLRVYVEELKPTPEGDLEILLQKWENGECAQKKIIAEKTKIPAVFKIDALNENKVLVLDTDYKKYLLFCMENSAEPEQSLACQCLVRTPEVDDEALEKFDKALKALPMHIRLSFNPTQLEEQCHI

**Lysozyme HEW (129 aa):**

KVFGRCELAAAMKRHGLDNYRGYSLGNWVCAAKFESNFNTQATNRNTDGSTDYGILQINSRWWCNDGRTPGSRNLCNIPCSALLSSDITASVNCAKKIVSDGNGMNAWVAWRNRCKGTDVQAWIRGCRL

**Myoglobin SpermWhale (153 aa):**

VLSEGEWQLVLHVWAKVEADVAGHGQDILIRLFKSHPETLEKFDRFKHLKTEAEMKASEDLKKHGVTVLTALGGILKKKGHHEAELKPLAQSHATKHKIPIKYLEFISDAIIHVLHSRHPGDFGADAQGAMNKALELFRKDIAAKYKELGYQG

**RNaseA Bovine (124 aa):** KETAAAKFERQHMDSSTSAASSSNYCNQMMKSRNLTKDRCKPVNTFVHESLADVQAVCSQKNVACKNGQTNCYQSYSTMSITDCRETGSSKYPNCAYKTTQANKHIIVACEGNPYVPVHFDASV

**alpha Lactalbumin Bovine (123 aa):**

EQLTKCEVFRELKDLKGYGGVSLPEWVCTTFHTSGYDTQAIVQNNDSTEYGLFQINNKIWCKDDQNPHSSNICNISCDKFLDDDLTDDIMCVKKILDKVGINYWLAHKALCSEKLDQWLCEKL

**Ribonuclease Bovine (124 aa):**

KETAAAKFERQHMDSSTSAASSSNYCNQMMKSRNLTKDRCKPVNTFVHESLADVQAVCSQKNVACKNGQTNCYQSYSTMSITDCRETGSSKYPNCAYKTTQANKHIIVACEGNPYVPVHFDASV

**Variable Positions in the TM Core and their contribution to β_s**

**Table S7.**
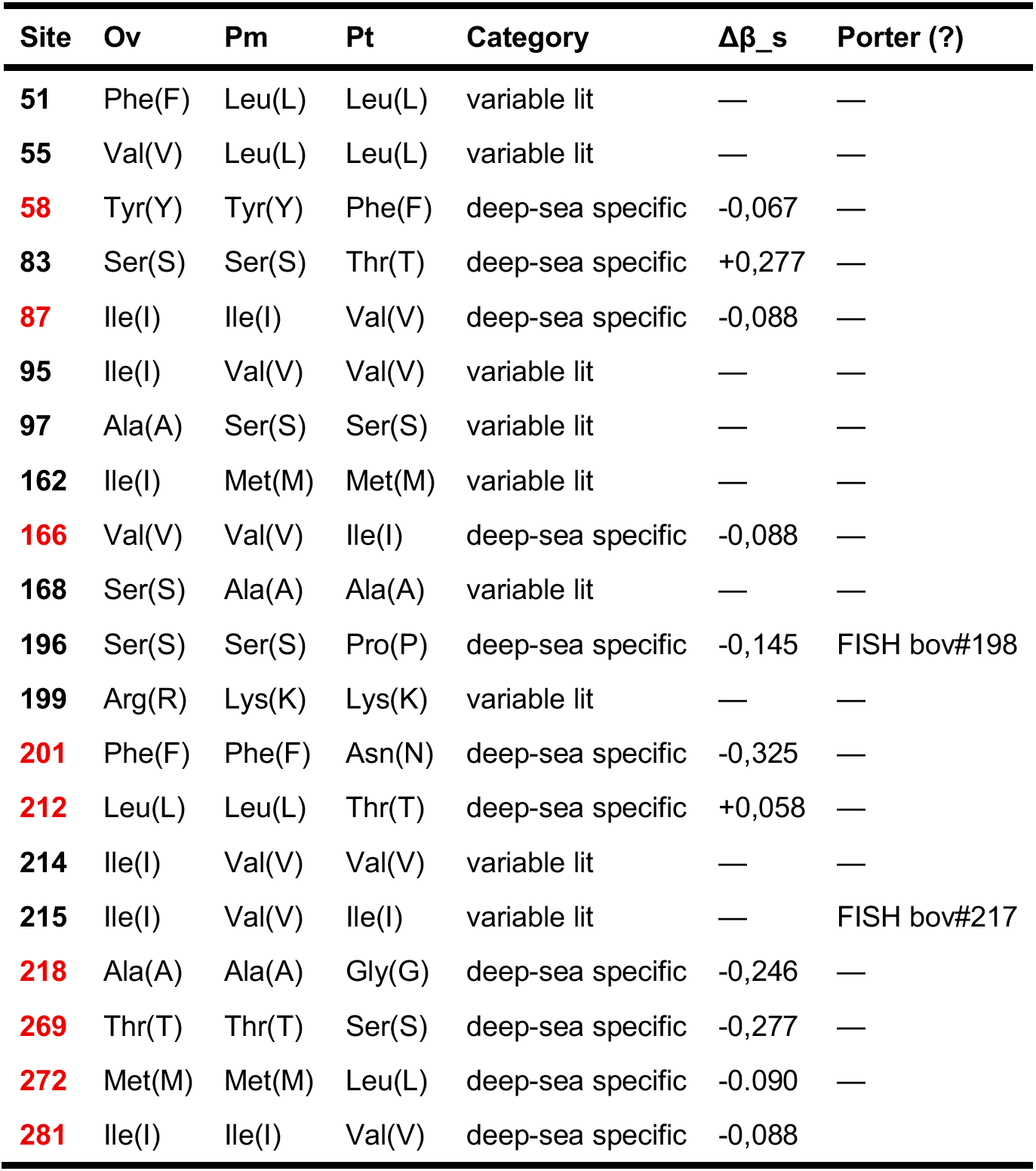
Amino acid differences in r-opsin1 among *O. vulgaris* (Ov), *P. macropus* (Pm) and *P. tetracirrhus* (Pt), focusing on the contribution of Pt-specific substitutions to Δβ_s, and highlighting positions with potential homology (?) to sites identified as under selection in Porter et al. (2016).

